# Chimeric Antigen Receptors Discriminate Between Tau and Distinct Amyloid-Beta Species

**DOI:** 10.1101/2025.02.05.636350

**Authors:** Cynthia J. Siebrand, Nicholas J. Bergo, Suckwon Lee, Julie K. Andersen, Chaska C. Walton

## Abstract

The lack of a definitive cure for Alzheimer’s disease (AD) is fueling the search for innovative therapeutic strategies. Having revolutionized cancer immunotherapy, immune cell engineering with chimeric antigen receptors (CAR) is being explored to target AD. Whether CARs can recognize distinct amyloid-β (Aβ) species and tau neurofibrillary tangles (NFTs)—hallmark pathologies of AD—remains unclear. To investigate this, we engineered CARs based on AD antibodies targeting tau (E2814), Aβ (Lecanemab and Aducanumab), and truncated pyroglutamate form of Aβ (Aβp3-42; Donanemab and Remternetug). To evaluate CAR function, we established the murine DO11.10 hybridoma T-cell line as a practical and scalable testing platform. Our findings demonstrate that CARs can detect and discriminate between tau preformed fibrils (PFFs), Aβ_1-42_, and Aβp3-42 aggregates. This highlights the potential of repurposing AD antibodies for CAR-based therapies to selectively target tau NFTs and distinct forms of Aβ senile plaques.

## Introduction

Recent advancements in Aβ-targeting antibodies have marked a significant milestone in AD research^1^. These therapies have been demonstrated to effectively reduce Aβ aggregates, a major hallmark of the disease^2–4^. Despite advancements, a definitive cure for AD is still lacking, and research continues to explore new therapeutic avenues^5^. A novel strategy leverages engineered immune cells^6–8^, which have been among the most impactful innovations in cancer treatment over the past decade^9^. Although many other synthetic receptor types are being actively explored^10^, clinical cell-based cancer interventions primarily consist of cytotoxic T lymphocytes engineered with CARs targeting tumor antigens^9^. CARs combine the antigen specificity of antibodies with T-cell receptor (TCR) signaling and the effector functions of T cells, enabling directed cell responses^11^.

The effector function of a specific CAR depends on the type of cell on which it is expressed. In treating hematologic malignancies, CARs expressed on cytotoxic T lymphocytes (CAR-T) specifically direct the cytotoxic function of the immune cells toward tumor cells^9,11^. Conversely, when CARs are expressed on regulatory T cells (Tregs), they redirect the immunomodulatory/anti-inflammatory function of these CAR-Tregs to protect against autoimmunity, prevent transplant rejection, and ameliorate neurodegenerative pathologies^12^. In the context of AD, Aβ-targeting CAR-Tregs have exhibited immunosuppressive functions *in vitro*^6^. This approach is promising as Aβ-specific TCR-engineered Tregs have been shown to attenuate AD pathobiology in APP/PS1 mouse models^7^. Recently, CARs have also been tested in cells other than T cells. Using the phagocytic common γ chain of the Fc receptor (FcRγ) as the intracellular signaling domain instead of TCR signaling, CAR-expressing macrophages (CAR-Ms) have demonstrated targeted clearing of Aβ plaques in APP/PS1 transgenic mouse^8^.

The region responsible for the CAR’s precise targeting ability is the antigen-binding domain, which often consists of a single-chain variable fragment (scFv) derived from antibodies with known specificities^11^. This strategy has been foundational for their use in cancer treatments, where CAR-T cell therapies such as the Food and Drug Administration (FDA)-approved CAR-T products Kymriah and Yescarta, utilize scFvs derived from antibodies to target CD19 in B-cell malignancies^13,14^. This strategy has also been capitalized in AD, in which an scFv based on the Aβ-targeting antibody Aducanumab guided the aforementioned CAR-M to Aβ plaques^8^.

Deriving the scFv from antibodies to enable precise targeting is not limited to CARs. Other synthetic receptors, such as the synthetic Notch (synNotch) receptor, can be engineered into immune cells to synthesize and secrete therapeutic antibodies when they bind to their targets^15,16^. Our lab recently demonstrated that synNotch containing scFvs based on Aducanumab (Aduhelm®) can detect Aβ_1-42_ preparations enriched in oligomers and deliver chimeric human-mouse versions of FDA-approved Lecanemab (Leqembi®) and Aducanumab^17^. Further, recent work *in vivo* demonstrated that synNotch receptors can deliver therapeutic payloads to the brain, targeting cancer or neuroinflammatory processes^18^. These studies highlight the importance of identifying functional scFvs as an innovative strategy for targeting engineered cells to the central nervous system to treat disorders, including AD.

Numerous antibodies have been extensively characterized for AD, with well-documented affinities for various protein species and subspecies implicated in key neuropathological hallmarks of AD^1,19^. Theoretically, this breadth of selectivity can be leveraged by scFv in engineered cells to strategically target pathological features in AD. For example, Braak stages establish AD severity based on the neuroanatomical spread of tau pathology^20,21^. Consequently, a tau-targeting CAR could concentrate engineered immune cells at these neuroanatomical regions to prevent the spread of tau pathology. Thus, the extensive knowledge base on existing AD antibodies provides an excellent foundation for synthetic receptor biology. However, the extent to which the specificity of these antibodies translates to scFv in CARs and other receptors is unknown.

To answer this, we developed second-generation CARs, incorporating scFvs derived from the investigational tau-targeting antibody E2814, the Aβ-targeting antibodies Aducanumab and Lecanemab, and the Aβp3-42-targeting antibodies Donanemab (Kisunla™) and Remternetug. For efficient and streamlined testing, these CARs were expressed in DO11.10 cells, a mouse CD4+ hybridoma T-cell line with monoclonal TCRs specific for the 329–337 epitope in the chicken ovalbumin 323–339 peptide (OVA)^22^. Our results demonstrate that DO11.10 cells are an effective and user-friendly platform for evaluating mouse CAR constructs. Notably, DO11.10 cells expressing the tau-targeting E2814-CAR, the Aβ-targeting Adu-CAR, or the Aβp3-42-targeting Don-CAR strongly responded to their cognate targets. These findings confirm that readily available scFv sequences can be used effectively to direct engineered immune cells toward specific pathological features of AD, providing a valuable approach for precision immunotherapy development.

## Results

### Engineering and evaluation of AD-targeting antibody-based CAR designs

To create CAR constructs for AD interventions, a second-generation CAR sequence from the murine CD19-targeting 1D3-28Z CAR served as the backbone (Fig. 1a; Table 1a)^23^. This CAR contains a CD28 costimulatory domain with a dileucine motif replaced by diglycine to enhance expression^24^. The antigen-binding domain sequence of 1D3-28Z was replaced with scFvs derived from the variable heavy chain (VHC) and variable light chain (VLC) of the following antibodies: tau-targeting antibody E2814 (E2814-CAR; Table 1b), Aβ-targeting antibodies Lecanemab (Lec-CAR; Table 1c) and Aducanumab (Adu-CAR; Table 1d), or the truncated pyroglutamate form of Aβ (Aβp3-42)-targeting antibodies Donanemab (Don-CAR; Table 1e) and Remternetug (Rem-CAR; Table 1f). All scFvs were constructed with the VLC followed by the VHC, linked by a G4S(3) flexible linker (Fig. 1a). The constructs were transduced into DO11.10 hybridoma T cells using a Murine Stem Cell Virus (MSCV) gamma retroviral vector (Fig. 1b). These cells possess functional TCR signaling machinery^22^, enabling the assessment of cell activation mediated by CARs and their CD3ζ signaling domain. The expression cassette included an enhanced green fluorescent protein (EGFP) reporter gene (Fig. 1b) for fluorescence-activated cell sorting (FACS) of transduced cells (Fig. 1c) and subsequent purification after clonal expansion for downstream analyses (Fig. 1d).

**Figure 1.**
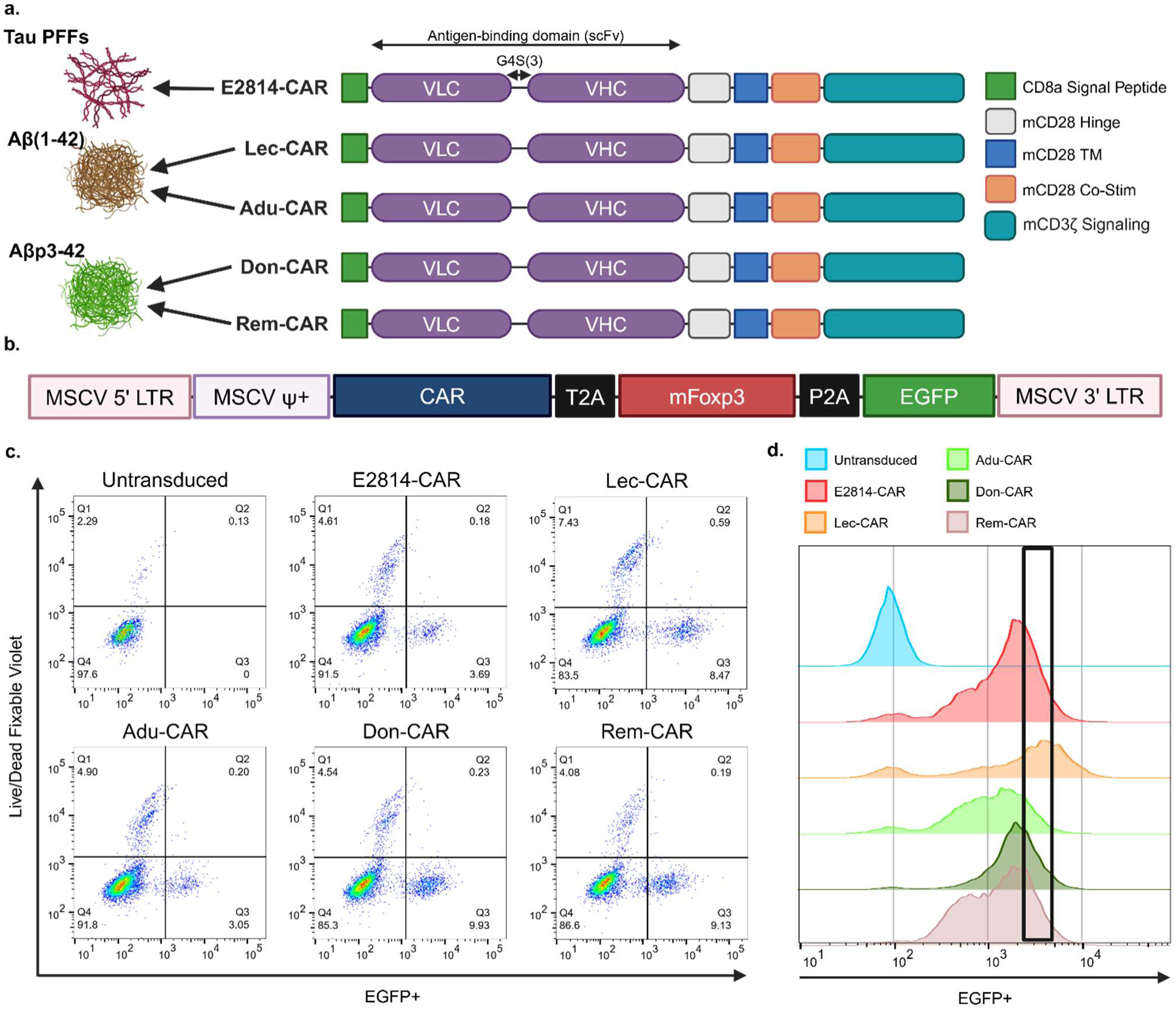
Engineering and evaluation of CAR constructs based on Alzheimer’s Disease-targeting antibodies. (**a**) Detailed schematic of the CAR constructs targeting tau PFFs, Aβ_1-42_, and Aβp3-42. Each construct features a human CD8α signal peptide (green) followed by an antigen-binding domain consisting of an scFv with the VLC (purple) followed by the VHC (purple) from antibodies E2814 (E2814-CAR), Lecanemab (Lec-CAR), Aducanumab (Adu-CAR), Donanemab (Don-CAR), or Remternetug (Rem-CAR), linked via a G4S(3) flexible linker. The scFv is fused to a murine CD28-based hinge (grey), transmembrane (TM) domain (blue), costimulatory domain (orange), and a mouse CD3ζ signaling domain (teal). Component sequences for each CAR construct are detailed in Table 1a–f. (**b**) Overview of the multicistronic retroviral vector system used for the experiments containing an MSCV backbone, the CAR construct, a murine Foxp3 (mFoxp3), and EGFP reporter gene, separated by T2A and P2A sequences. An assessment of the Foxp3 is not incorporated in the current study (see Materials and Methods for additional information) (**c**) Representative flow cytometry plots of EGFP and viability dye staining (Live/Dead Fixable Violet) used to sort transduced cell clones (Q3) via FACS. (**d**) Subsequent FACS-based selection strategy after expansion using a standardized EGFP signal intensity range (black region) to ensure homogeneous CAR-expression in DO11.10 cell clones used for the experiments.

### Tau PFFs 3.125-200 nM do not significantly affect cell viability

Tau aggregates, along with Aβ, represent the key neuropathologies of AD^2–4^. Moreover, tau pathology serves as the basis for Braak staging of AD^20,21^. Tau tangles are believed to seed and propagate the spread of tau pathology and the tau-targeting investigational antibody E2814 aims to prevent this propagation^25^. E2814 is a monoclonal IgG1 antibody, which binds to an epitope within the tau protein’s microtubule-binding domain near the mid-region—an essential site involved in seeding and propagating pathological tau aggregates. In this study, we developed the E2814-CAR (Fig. 1a, Table 1b) to evaluate whether CARs can target pathological tau aggregates. As controls, we used untransduced DO11.10 cells and the Aβ-targeting Lec-CAR (Fig. 1a, Table 1c), which is based on the FDA-approved Lecanemab. To assess the ability of each CAR construct to detect tau, we measured activation by CD69 immunostaining^26^ and IL-2 secretion^27^ using flow cytometry and enzyme-linked immunosorbent assay (ELISA), respectively (Fig. 2a).

**Figure 2.**
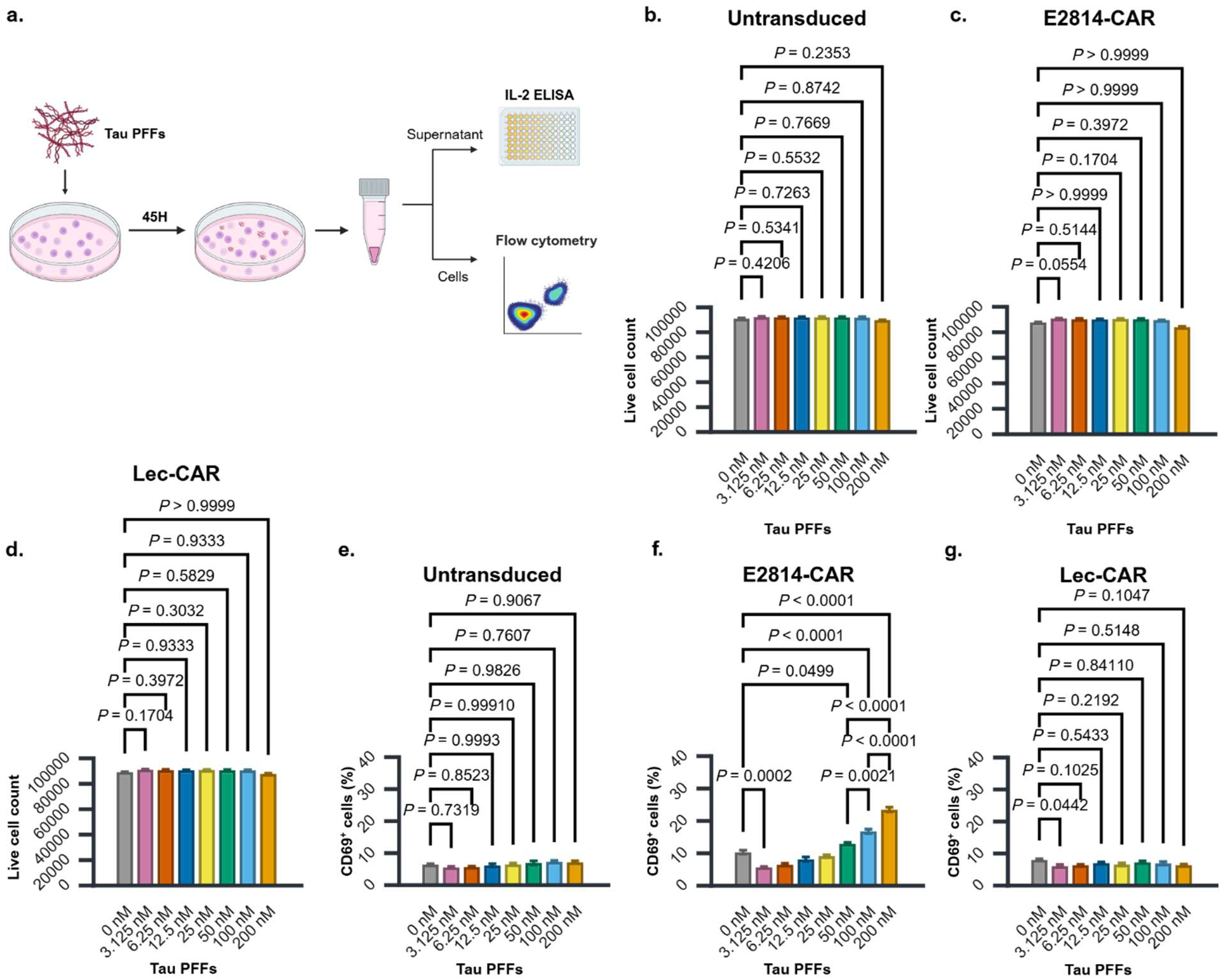
Tau PFFs selectively activate E2814-CAR-expressing cell clones without inducing cell death. (**a**) Schematic overview of the experiments. After tau PFFs treatments for 45 hours, the media was collected for analysis of IL-2 by ELISA and the cells were harvested for analysis of EGFP expression, viability, and CD69 expression by flow cytometry. (**b-d**). Total number of live cells remaining after tau PFFs treatment for (**b**) untransduced, (**c**) E2814-CAR, and (**d**) Lec-CAR clones. (**e-g**) Percentage of CD69-positive (CD69^+^) live cells for (**e**) untransduced, (**f**) E2814-CAR, and (**g**) Lec-CAR DO11.10 cell clones. Only relevant statistical p-values are shown; full statistical analyses available in Tables 2a-c & Tables 3a-c. Statistical comparisons: (**b**) Dunnett, (**c**, **d**) Dunn’s, and (**e-g**) Tukey multiple comparisons. Three biological replicates. Statistical significance declared at *p* < 0.05. Graphs display mean ± s.e.m.

High percentages of live EGFP^+^ cells, ranging from 95.56% to 98.00%, were observed in E2814-CAR and Lec-CAR (Sup. Fig. 1a, b), confirming active and homogeneous CAR expression across clones. To assess the impact of tau PFF treatment on cell viability, we analyzed the total number of viable cells across different experimental conditions. Statistically significant differences were detected among treatment conditions for untransduced cells (*F*(7, 16) = 3.0883, *p* = 0.0293, *η²* = 0.5747; Fig. 2b; Table 2a). To avoid overcorrection of the family wise error, we performed Dunnett multiple comparisons between tau PFFs treatments and the reference vehicle-control (VEH) group only. The analysis revealed that there were no significant differences between the VEH and tau PFFs treatments (Fig. 2b), indicating that the differences detected by the omnibus test were likely within tau PFFs treatments. For CAR-expressing cells, significant differences were detected for E2814-CAR (*x^2^*(7) = 17.1200, *p* = 0.0166; Fig. 2c; Table 2b) and Lec-CAR clones (*x^2^*(7) = 14.1200, *p* = 0.0491; Fig. 2d; Table 2c). However, multiple comparisons of treatment concentrations against VEH revealed no significant differences for E2814-CAR (*p* ≥ 0.0554; Fig. 2c) or Lec-CAR (*p* ≥ 0.1704; Fig. 2d), suggesting that omnibus statistical differences were also likely due to variability within 3.125–200 nM treatment conditions. The bar graphs reflecting viability based on the percentage of live cells were also consistent with a lack of significant cytotoxicity (Sup. Fig. 1c, d).

### Tau PFFs increased the percentage of CD69-positive E2814-CAR-expressing cells

To determine whether the CAR-expressing cells were activated in response to tau PFF treatments, we assessed the expression of the T-cell activation marker CD69 in untransduced (Fig. 2e; Sup. Fig. 2), E2814-CAR (Fig. 2f; Sup. Fig. 2), and Lec-CAR (Fig. 2g; Sup. Fig. 2) expressing DO11.10 cells. For untransduced cells, statistical analysis revealed significant differences among tau PFF treatment groups (*F*(7, 16) = 2.7528, *p* = 0.0442, *η²* = 0.5464; Table 3a). However, Tukey all-pairwise comparisons tests did not detect any statistical differences (Fig. 2e). The closest comparison to significance was between 3.125 and 100 nM treatments (*p* = 0.0823; Table 3a). The discrepancy between the omnibus test and pairwise comparisons likely reflect Tukey’s correction for family-wise error. Importantly, relative to VEH, none of the tau PFFs treatments in untransduced cells displayed values close to statistical significance (*p* ≥ 0.7319; Fig 2e), evidencing the lack of activation of untransduced DO11.10 cells.

Similarly, in the Lec-CAR group a non-significant trend was observed (*F*(7, 16) = 2.5724, *p* = 0.0557, *η²* = 0.5295; Table 3c). However, no tau PFF treatments resulted in a statistically significant increase in CD69-positive relative to VEH (Fig. 2g). Instead, there was only a decrease for tau PFFs at 3.125 nM (*p* = 0.0442; Fig. 2g). In contrast, cells expressing E2814-CAR exhibited highly significant differences (*F*(7, 16) = 127.2703, *p* < 0.0001, *η²* = 0.9824; Table 3b). Compared to VEH, the percentage of CD69-positive cells was significantly increased at 50 (*p* = 0.0499), 100 (*p* < 0.0001), and 200 nM (*p* < 0.0001) concentrations (Fig. 2f). The increase was also significant from 50 to 100 nM (*p* = 0.0021) and from 100 to 200 nM (*p* < 0.0001) tau PFFs concentrations, demonstrating a concentration-dependent response of E2814-CAR to tau PFFs.

### Tau PFFs induce IL-2 secretion in cells expressing E2814-CARs

The percentage of CD69-positive cells represents a snapshot of activation at the assay’s endpoint. To gain insight into activation dynamics and the magnitude of the response throughout the assay, IL-2 levels accumulated in the media were measured via ELISA. IL-2 levels in untransduced cells were below the 3.1 pg/mL detection threshold (orange line) of the ELISA kit specified by the vendor (Fig. 3a), indicating no response to the treatments and were excluded from further analysis. Interestingly, VEH-treated CAR-expressing cells exhibited significantly higher IL-2 levels compared to untransduced cells (Fig. 3a), consistent with the known ligand-independent tonic signaling observed in CARs^28^. For Lec-CAR-expressing cells, IL-2 production decreased across all tau PFF concentrations relative to VEH (black dotted lines; Fig. 3a), aligning with CD69 analysis (Fig. 2g). Due to a patent lack of response, IL-2 production by Lec-CAR was not further analyzed. For E2814-CAR clones, only tau PFFs treatments at 200 nM were clearly above VEH (Fig. 3a), and statistical analysis showed this difference was significant (*t*(4) = −4.4716, *p* = 0.0111; Fig. 3b; Table 4), consistent with CD69-based activation assessments (Fig. 2f).

**Figure 3.**
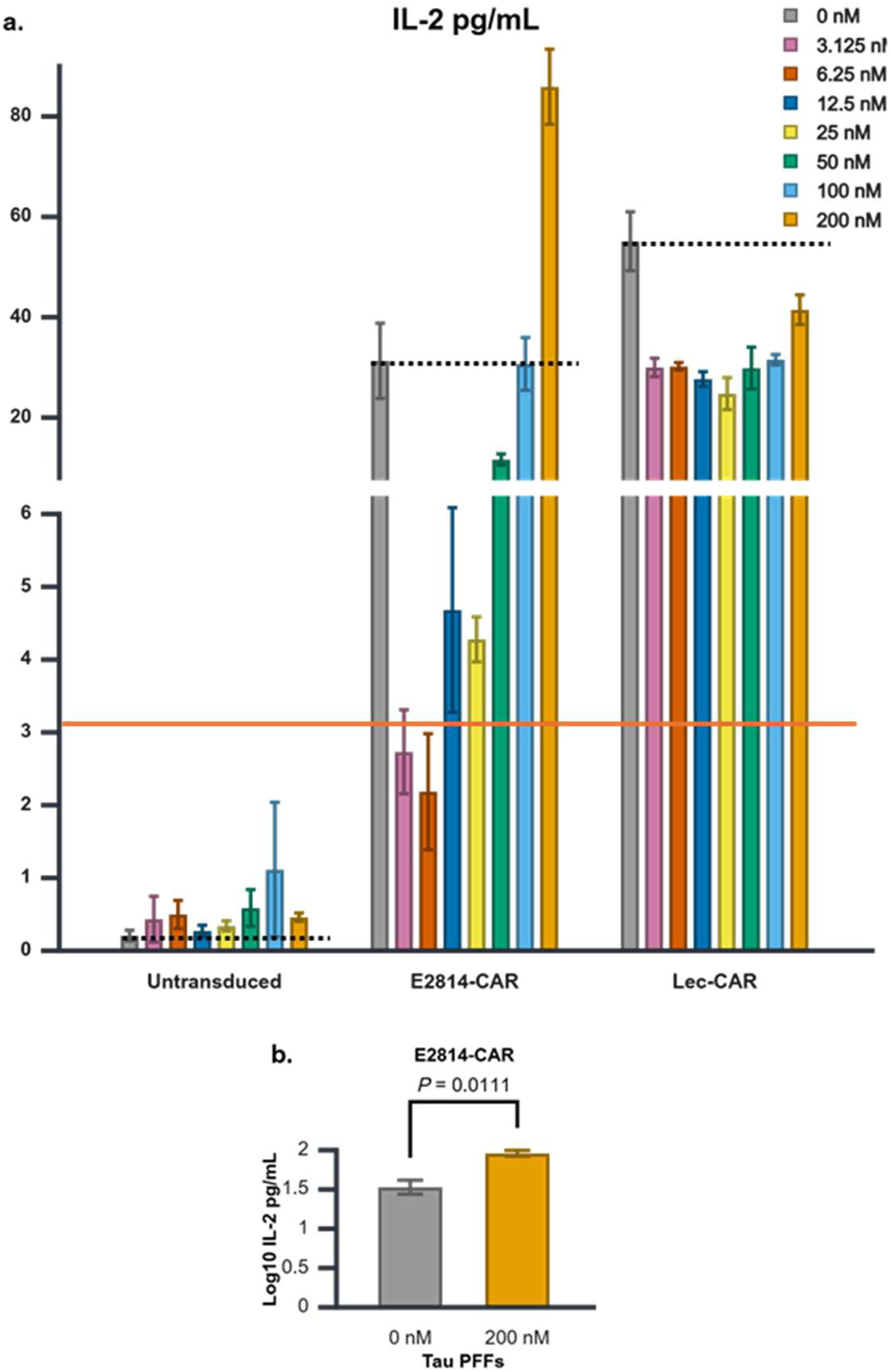
E2814-CAR-expressing cell clones secrete IL-2 in response to tau PFF treatment. (**a**) Bar chart of IL-2 concentrations measured in the media of tau PFF-treated CAR clones shown for two concentration ranges: 0–6.2 pg/mL (bottom) and 8–90 pg/mL (top). Note that each range is displayed on a different scale. IL-2 levels in untransduced cells at all doses and in E2814-CAR at 3.125 and 6.25 nM are below the assay’s 3.1 pg/mL detection threshold (orange line). E2814-CAR IL-2 levels remain below 0 nM (VEH; black dotted lines) at 3.125–100 nM tau PFFs, while Lec-CAR-expressing cells show no IL-2 increase at any concentration. (**b**) Statistical analysis comparing IL-2 production at 0 nM (VEH) and 200 nM tau PFFs in E2814-CAR-expressing cells shows a significant increase. Statistical test: independent two-tailed *t*-test. Three biological replicates. Statistical significance declared at *p* < 0.05. Graphs display mean ± s.e.m.

To summarize tau PFFs experiments, tonic CAR activation was evidenced by the increased activation in VEH-treated CAR clones relative to VEH-treated untransduced cells (Fig. 2e-g; Fig. 3a). The reduction in IL-2 production (Fig. 3a) and the percentage of CD69-positive cells (Fig. 2g) observed for Lec-CAR relative to VEH likely reflects a nonspecific tau PFF-induced suppression of tonic activation, an effect that also impacted E2814-CAR clones (Fig. 3a; Fig. 2f). As a result, for a specific E2814-CAR response to tau PFFs to be detected, it would need to overcome this nonspecific suppression. Consistent with this, E2814-CAR clones treated with low tau PFF concentrations exhibited a marked reduction in IL-2 levels (Fig. 3a) and percentage of CD69-positive cells (Fig. 2f) relative to VEH. However, with increasing tau PFF concentrations, activation metrics progressively escalated, indicating that specific CAR activation was overcoming suppression, ultimately leading to activation that significantly surpassed VEH levels (Fig. 2f; Fig. 3b).

### Aβ_1-42_ shows varying cytotoxic effects on CAR-expressing cells

To evaluate the ability of CAR constructs to detect Aβ_1-42_ preparations enriched in oligomers, we utilized Lec-CAR (Fig. 1a; Table 1c) and Adu-CAR (Fig. 1a; Table 1d). Aducanumab is a human IgG1 monoclonal antibody isolated from cognitively normal healthy aged individuals with selectivity for aggregated Aβ oligomers over monomers and Lecanemab is a humanized IgG1 monoclonal antibody with selectivity for soluble Aβ oligomers and protofibrils over monomers and fibrils^1,29–33^. As controls, we used untransduced cells and the tau-targeting E2814-CAR (Fig. 1a; Table 1b). To assess CAR function, the cells were exposed to different concentrations of aggregated Aβ_1-42_ preparations enriched in oligomers for 48 hours (Fig. 4a) after which EGFP expression, viability, CD69 expression, and IL-2 secretion were measured.

**Figure 4.**
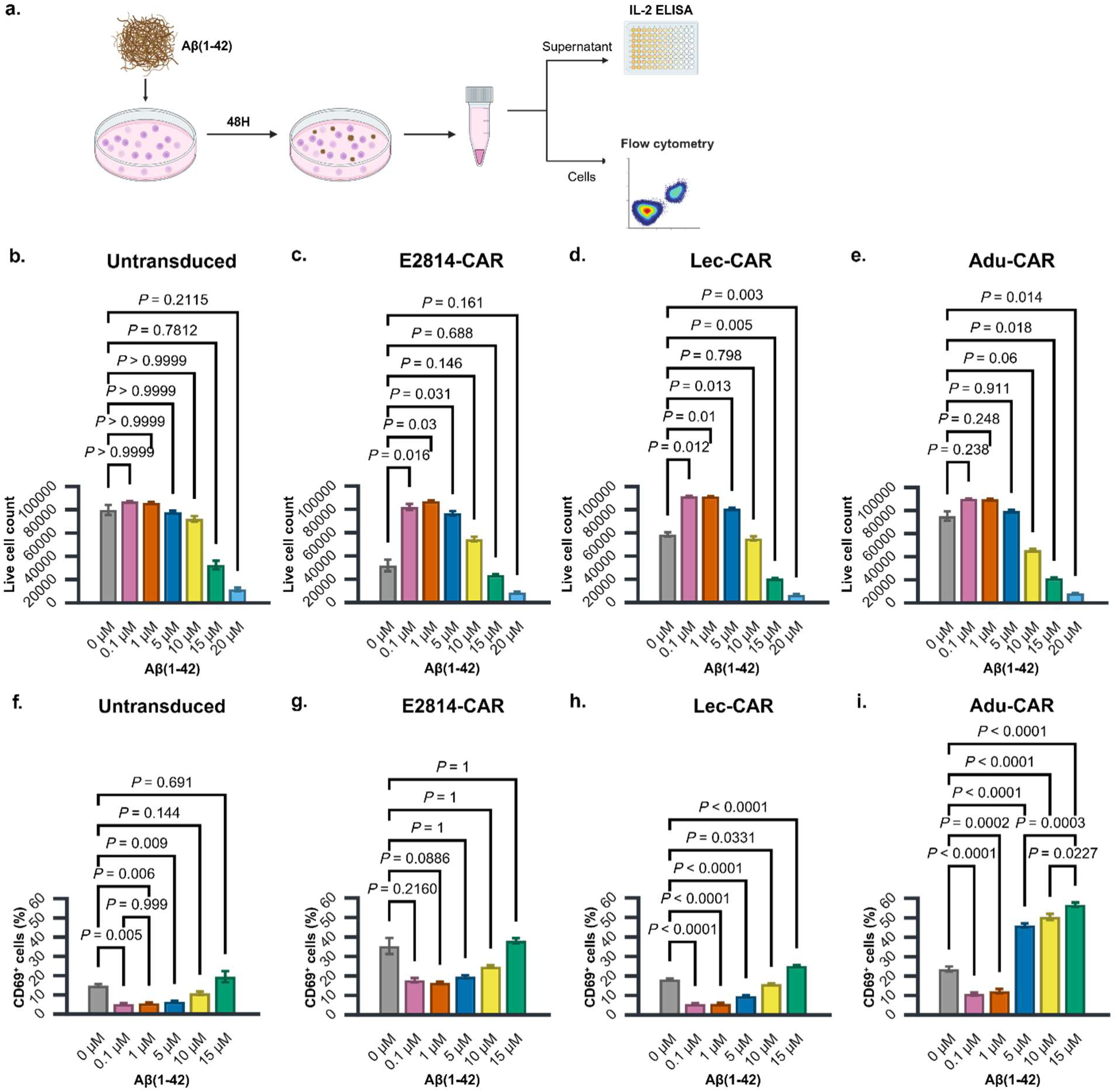
Cytotoxic Aβ_1-42_ treatments selectively activate Adu-CAR and Lec-CAR-expressing cell clones. (**a**) Schematic overview of the experiments. After Aβ_1-42_ treatments for 48 hours, the media was collected for analysis of IL-2 by ELISA and the cells were harvested for analysis of EGFP expression, viability, and CD69 expression by flow cytometry. (**b-e**) Total number of live cells remaining after Aβ_1-42_ treatment for (**b**) untransduced, (**c**) E2814-CAR, (**d**) Lec-CAR, and (**e**) Adu-CAR clones. (**f-i**) Percentage of CD69-positive (CD69^+^) live cells analyzed for (**f**) untransduced, (**g**) E2814-CAR, (**h**) Lec-CAR, and (**i**) Adu-CAR clones. Statistical comparisons: (**b, g**) Dunn’s, (**c-f**) Games-Howell, and **(h, i**) Tukey multiple comparisons. Only relevant statistical *p*-values are shown; full statistical analyses available in Tables 5a,c-e & Tables 6a-d. Three biological replicates. Statistical significance declared at *p* < 0.05. Graphs display mean ± s.e.m.

High percentages of live EGFP+ cells, ranging from 89.46% to 95.00%, were observed in E2814-CAR, Lec-CAR, and Adu-CAR (Sup. Fig. 3a, b), confirming active and homogeneous CAR expression across clones. However, in contrast to tau PFF viability assessment, Aβ_1-42_ treatments revealed cytotoxic effects. Visual inspection of the bar graphs indicated a reduction in the total number of untransduced cells at higher Aβ_1-42_ concentrations (Fig. 4b). Assumption testing revealed that the Aβ_1-42_ 15 μM treatment group was not normally distributed (*p* = 0.0159; Table 5a). A Kruskal-Wallis nonparametric test revealed a significant effect of Aβ_1-42_ treatment (*x^2^*(6) = 18.3377, *p* = 0.0054; Table 5a), but post hoc analysis did not detect significant differences between ranks between any Aβ_1-42_ treatment condition and VEH (Fig. 4b). Given the reduced statistical power of non-parametric tests and the evident differences seen in the bar graphs (Fig. 4b), an additional parametric analysis was conducted by excluding the non-normally distributed 15 μM Aβ_1-42_ treatment group. This analysis revealed highly significant effects of Aβ_1-42_ treatment on viability (*F*(5,5.1765) = 351.3600, *p* < 0.0001; Table 5b), with multiple comparisons evidencing a decrease in viability at 20 μM Aβ_1-42_ (*p* = 0.007; Sup. Fig. 3c).

The effects of Aβ_1-42_ treatment on E2814-CAR viability were significant (*F*(6, 6.0292) = 1094.1400, *p* < 0.0001; Table 5c), but these treatments were attributed to a paradoxical increase in viability at lower concentrations (Fig. 4c;). A statistically significant increase in viability was also seen for Lec-CAR clones at 0.1-5 μM concentrations (*p* ≤ 0.013), but this was also accompanied by a significant decrease in viability at 15 (*p* = 0.0049) and 20 μM (*p* = 0.003) treatments (*F*(6, 5.7061) = 4636.5500, *p* < 0.0001; Fig. 4d; Table 5d). Adu-CAR clones’ loss of viability was also significant at 15 (*p* = 0.018) and 20 μM (*p* = 0.014) concentrations (*F*(6, 6.0781) = 5039.7700, *p* < 0.0001; Fig. 4e; Table 5e).

Interestingly, all groups showed a consistent trend of the total number of viable cells increasing at Aβ_1-42_ concentrations followed by decreases at high concentrations (Fig. 4b–e), which was consistent with viability as reflected by the percentage of live cells (Sup. Fig. 3d, e). Reduced viability aligns with the established toxic effects of Aβ^34^, while increased proliferation may involve activation of endogenous receptors for which Aβ is a known ligand^35^.

### Aβ_1-42_ increases the percentage of CD69-positive Adu-CAR and Lec-CAR-expressing cells

The results described above were corroborated by the assessment of the total number of viable CD69-positive cells, showing a reduction in activated cells at high Aβ_1-42_ concentrations (Sup. Fig. 4a). Interestingly, 20 μM Aβ_1-42_ treatments decreased the total number of CD69-positive cells (Sup. Fig. 4a) and simultaneously increased the percentage of CD69-positive cells (Sup. Fig. 4b, c). Since this trend was also observed in untransduced cells, the activation of the surviving viable cells could not be attributed to the CAR and may instead be a nonspecific response to widespread cell death. To avoid conflating nonspecific activation with Aβ_1-42_-specific CAR activation, statistical analysis of CD69-positive percentages excluded 20 μM Aβ_1-42_ treatments.

At the remaining concentrations, the analysis of the percentage of CD69-positive populations for untransduced cells revealed a highly significant effect of Aβ_1-42_ treatment (*F*(5,5.4992) = 22.8000, *p* = 0.0013; Table 6a). This effect was characterized by a significant reduction in the percent of CD69-positive cells at 0.1-5 μM Aβ_1-42_ treatments (*p* ≤ 0.009; Fig. 4f), followed by a recovery to VEH levels at higher concentrations (Fig. 4f). E2814-CAR-expressing cells exhibited a similar pattern (*x^2^*(5) = 15.4795, *p* = 0.0085; Fig. 4g; Table 6b). However, there were no increases above VEH in either untransduced (Fig. 4f) or E2814-CAR-expressing cells (Fig. 4g).

In contrast, highly significant differences were observed for Lec-CAR (*F*(5,12) = 278.3576, *p* < 0.0001, *η²* = 0.9915; Fig. 4h; Table 6c) and Adu-CAR (*F*(5,12) = 310.1872, *p* < 0.0001, *η²* = 0.9923; Fig. 4i; Table 6d). These effects corresponded to statistically significant increases in the percentage of CD69-positive cells at 15 μM Aβ_1-42_ (*p* < 0.0001) for Lec-CAR-expressing cells (Fig. 4h) and starting at 5 μM Aβ_1-42_ (*p* < 0.0001) for Adu-CAR-expressing cells (Fig. 4i). For Adu-CAR, there was also a significant stepwise increase from 5 (*p* = 0.0003) and 10 (*p* = 0.0227) to 15 μM Aβ_1-42_ treatments (Fig. 4i). Significant reductions in the percentage of CD69-positive cells at lower treatment concentrations were also detected for Lec-CAR (Fig. 4h) and Adu-CAR (Fig. 4i), corroborating the presence of nonspecific suppression by Aβ_1-42_.

Similar to tau PFF treatments (Fig. 2e–g), statistically significant reductions relative to VEH in CD69-positive percentages were observed across all groups (Fig. 4f–i). However, unlike tau treatments, reductions in activation with Aβ_1-42_ also affected untransduced cells (Fig. 4f). This suggests that Aβ_1-42_ has a broader suppressive effect than tau, reducing both the basal activation of CAR receptors and the overall activation of DO11.10 cells. Another notable phenomenon was a pattern of increased viability at lower Aβ_1-42_ concentrations (Fig. 4b–e), significant for some constructs (Fig. 4c, d). The significant reduction in CD69 expression at overlapping concentrations suggests that the enhanced viability is independent of DO11.10 cell activation (Fig. 4f–i).

### Aβ_1-42_ induces IL-2 secretion by cells expressing Adu-CARs

IL-2 levels in untransduced cells remained below the ELISA detection threshold (orange line; Fig. 5a). This was also true for 20 μM Aβ_1-42_ treatments, seemingly contradicting the apparent increase in CD69-positive cells at this concentration (Sup. Fig. 4b). However, the percentage of CD69-positive cells does not capture the magnitude of activation, which were likely orders of magnitude lower for untransduced cells. Under VEH conditions, IL-2 levels were 0.365 (SD 0.1796) pg/ml (Fig. 5a) and the number of activated CD69-positive cells was 11381.3333 (SD 1614.4635; Sup. Fig. 4a). For comparison, under VEH treatment, E2814-CAR clones produced 91.0603 (SD 4.7047) pg/ml (Fig. 5a) and had 9624 (SD 1153.8388) CD69-positive activated cell clones (Sup. Fig. 4a). Thus, despite having approximately 85% of the number of activated cells, tonically active E2814-CAR clones produced almost 250 times more IL-2. Moreover, while the percentage of CD69-positive untransduced cells at 20 μM Aβ_1-42_ (33.92%, SD 1.3787) was above VEH (14.39%, SD 1.3095; Sup. Fig. 4b), the total number of CD69-positive cells 3652.333 (SD 925.4503) was around 3 times less than VEH (Sup. Fig. 4a). Consequently, because of the weak response of untransduced cells, IL-2 levels below the ELISA kit detection threshold, even at the highest Aβ_1-42_ concentration, align with what can reasonably be expected.

**Figure 5.**
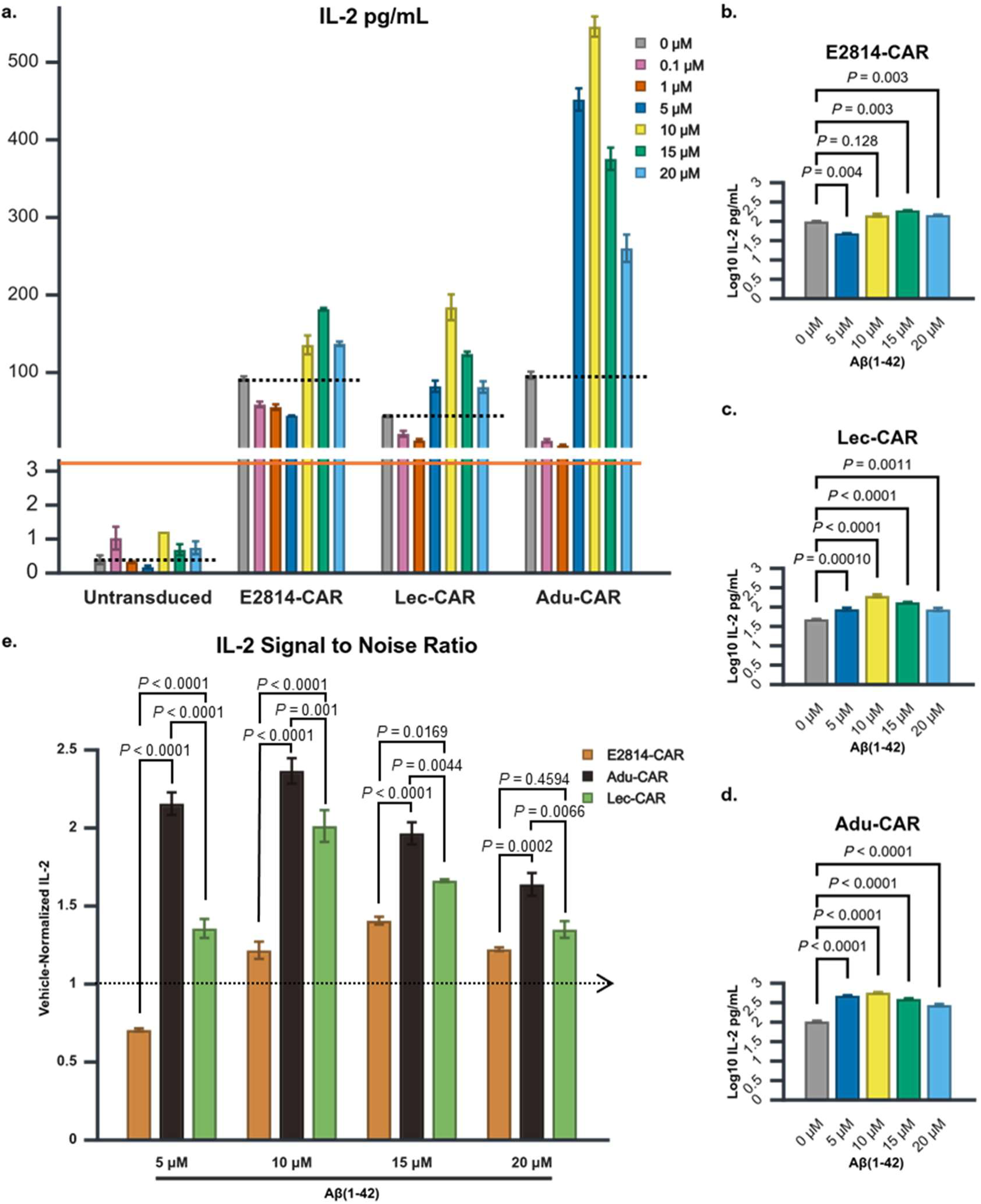
IL-2 secretion by Adu-CAR clones shows the strongest response to Aβ_1-42_ treatments. (**a**) Bar chart of IL-2 concentrations measured in the media of Aβ_1-42_-treated CAR clones displayed at two concentration ranges: 0–3.3 pg/mL (bottom) and 3.5–600 pg/mL (top). Note that each range is shown on a different scale. IL-2 levels in untransduced cells at all doses remain below the assay’s 3.1 pg/mL detection threshold (orange line). For E2814-CAR-expressing cells, IL-2 levels remain below 0 μM (VEH; black dotted lines) at 0.1–5 μM Aβ_1-42_. For Lec-CAR and Adu-CAR, IL-2 levels remain below VEH at 0.1 and 1 μM Aβ_1-42_. (**b-d**) Statistical analyses of IL-2 production in response to 0–20 μM Aβ_1-42_ for (**b**) E2814-CAR, (**c**) Lec-CAR, and (**d**) Adu-CAR. (**e**) Signal-to-noise ratio analysis comparing E2814-CAR, Lec-CAR, and Adu-CAR at each Aβ_1-42_ concentration. The black dotted arrow indicates VEH levels. Statistical comparisons: (**b**) Games-Howell, (**c, d**) Tukey, (**e**) and Bonferroni multiple comparisons. Only relevant statistical p-values are shown; full statistical analyses available in Tables 7a-d. Three biological replicates. Statistical significance was determined at *p* < 0.05. Graphs display mean ± s.e.m.

As per the rest of the CAR-expressing clones, none produced IL-2 levels above VEH at 0.1 and 1 μM Aβ_1-42_, prompting their exclusion from subsequent analysis. Omnibus tests for the reminder of the treatments revealed significant differences in IL-2 production for E2814-CAR (*F*(4,4.4205) = 3645.6600, *p* < 0.0001; Fig 5b; Table 7a), Lec-CAR (*F*(4,10) = 55.0529, *p* < 0.0001, *η²* = 0.9566; Fig 5c; Table 7b), and Adu-CAR DO11.10 cell clones (*F*(4,10) = 263.8516, *p* < 0.0001, *η²* = 0.9906; Fig 5d; Table 7c). While IL-2 levels in E2814-CAR-expressing cells exceeded VEH levels only at Aβ_1-42_ concentrations above 15 μM (*p* = 0.003; Fig. 5b), Lec-CAR (*p* ≤ 0.0011; Fig. 5c) and Adu-CAR (*p* < 0.0001; Fig. 5d) exhibited significant IL-2 increases at all tested Aβ_1-42_ concentrations.

These results suggest E2814-CAR responded to Aβ_1-42_ treatment (Fig. 5b), raising questions about the specificity of Lec-CAR (Fig. 5c) and Adu-CAR (Fig. 5d) clones. In fit, CD69 analysis in untransduced cells previously indicated Aβ_1-42_ can nonspecifically activate DO11.10 cells at 20 μM (Sup. Fig. 4b). Owed to the inherent differences in the assessment of activation (described in Sup. Fig. 5a, b), the analysis of IL-2 may capture nonspecific activation at different concentrations. If nonspecific activation is the only driving force for all CAR constructs, IL-2 production by E2814-CAR, Lec-CAR, and Adu-CAR-expressing cells should be similar. A direct comparison of IL-2 production between CAR clones was not possible because they had different levels of tonic activation (Fig. 5a). For example, E2814-CAR basally produced more IL-2 than Lec-CAR when treated with VEH (Fig. 5a), therefore having a higher baseline IL-2 production added to all Aβ_1-42_ treatments. This would confound any direct comparisons of Aβ_1-42_ treatments. To overcome this problem, we VEH-normalized IL-2 levels to obtain the signal-to-noise ratio, which allowed for the comparison of the fold-increase in IL-2 production over VEH between CAR clones (see Materials and Methods for additional details).

The analysis of the signal-to-noise ratio showed that Adu-CAR and Lec-CAR IL-2 production was above VEH (black dotted arrow) at Aβ_1-42_ treatment concentrations from 5-20 μM (Fig. 5e). For E2814-CAR, this was only true for concentrations of 10 μM Aβ_1-42_ and above. A two-way ANOVA revealed a significant interaction between CAR type and Aβ_1-42_ (*F*(6,24) = 18.3852, *p* < 0.0001, partial *η²* = 0.8213; Fig 5e; Table 7d). Bonferroni-corrected all pairwise comparisons indicated that the signal-to-noise ratio of Adu-CAR was significantly higher than Lec-CAR (*p* ≤ 0.0066) and E2814-CAR (*p* ≤ 0.0002; Fig. 5e) at all tested concentrations. This ratified the qualitatively stronger response of Adu-CAR that was already discernible by visual inspection of Fig. 5a. While Lec-CAR and E2814-CAR were within similar ranges of IL-2 production (Fig. 5a), Lec-CAR exhibited a significantly higher signal-to-noise ratio than E2814-CAR at all concentrations (*p* ≤ 0.0169), except at 20 μM Aβ_1-42_ (*p* = 0.4594; Fig. 5e). Thus, the signal-to-noise ratio for the Aβ-targeting CARs was superior to that of the tau-targeting CAR, indicating Aβ_1-42_-specific responses consistent with the analysis of CD69 (Fig. 4h-i).

In summary, the analysis of CAR response to Aβ_1-42_ treatments included effects that nonspecifically altered activation, which influenced metrics differently depending on whether it was assessed by CD69 expression (see Sup. Fig. 5a for details) or IL-2 production (see Sup. Fig. 5b for details). In this context, Adu-CAR clones were qualitatively superior to Lec-CAR clones, responding to a wider range of concentrations more strongly.

### Aβp3-42 induces IL-2 secretion by cells expressing Rem-CARs

To evaluate the ability of CAR constructs to detect Aβp3-42, we designed Don-CAR (Fig. 1a; Table 1e), based on Donanemab, and Rem-CAR (Fig. 1a; Table 1f), based on Remternetug. Donanemab is the third FDA-approved Aβ-targeting antibody for AD, while Remternetug is a follow-up of Donanemab currently in phase III clinical trials^1,5^. These antibodies are specifically designed to target insoluble Aβp3-42-rich plaques, particularly core plaques deposited in the brain parenchyma, while avoiding soluble Aβ species^1,36,37^. The response of each CAR construct to Aβp3-42 was measured using IL-2 secretion ELISA assays (Fig. 6a). Alongside the Aβp3-42-targeting CARs, untransduced cells and control CAR clones, including E2814-CAR and Lec-CAR, were also tested.

**Figure 6.**
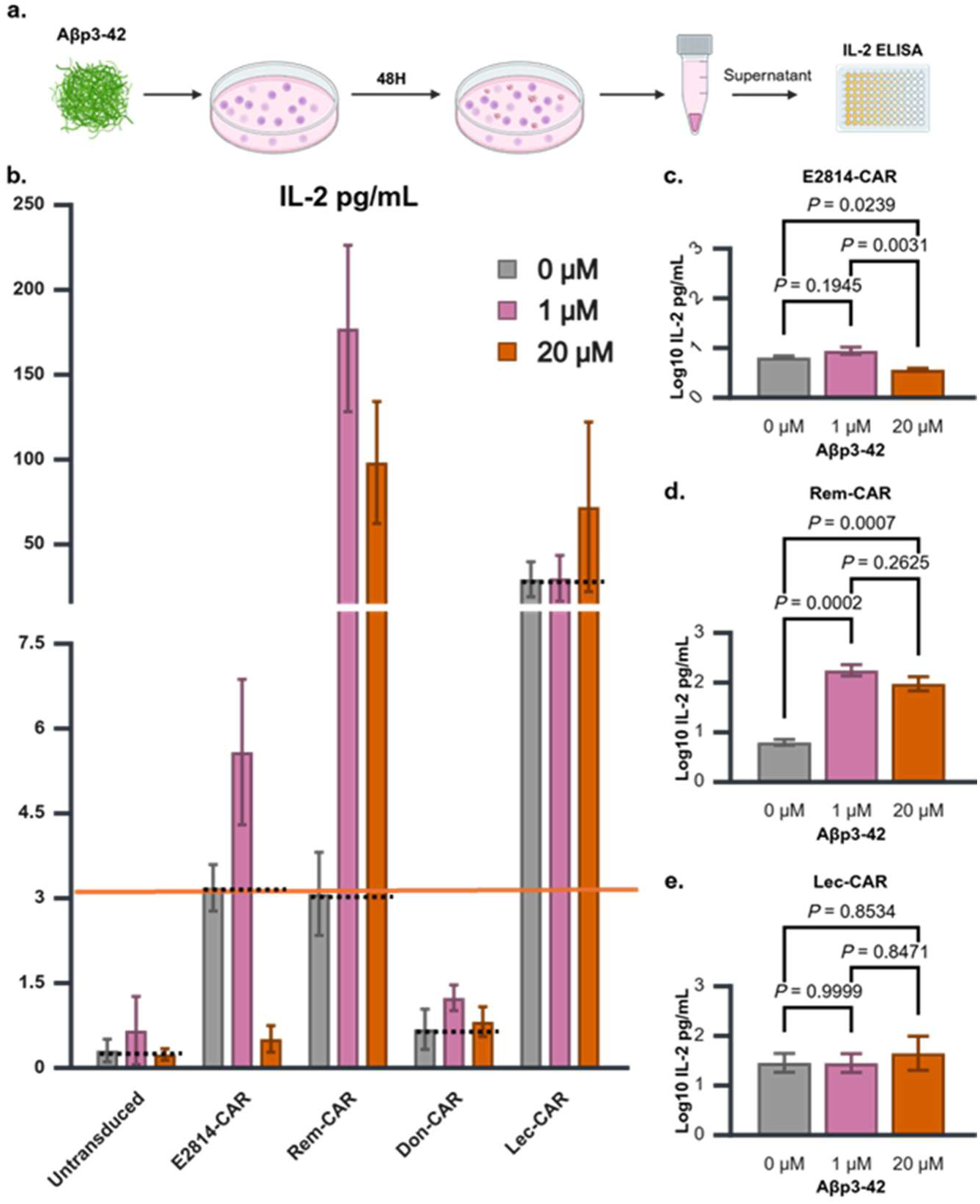
Rem-CAR-expressing cell clones secrete IL-2 in response to Aβp3-42 treatment. **(a)** Schematic overview of the experiments. After Aβp3-42 treatments for 48 hours, the media was collected for analysis of IL-2 by ELISA. (**b**) Bar chart of IL-2 concentrations measured in the media of Aβp3-42-treated CAR clones, displayed at two concentration ranges: 0–7.9 pg/mL (bottom) and 15–250 pg/mL (top). Each range is shown on a different scale. IL-2 levels in untransduced cells and Don-CAR clones at all doses, as well as E2814-CAR at 20 μM, are below the assay’s 3.1 pg/mL detection threshold (orange line). With the exception of E2814-CAR at 20 μM, IL-2 secretion in E2814-CAR, Lec-CAR, and Rem-CAR-expressing cells remains above 0 μM (VEH; black dotted line) across all concentrations. (**c-e**) Statistical analyses of IL-2 production in response to 0–20 μM Aβp3-42 for (**c**) E2814-CAR, (**d**) Rem-CAR, and (**e**) Lec-CAR. Statistical comparisons: (**c-e**) Tukey multiple comparisons. Only relevant statistical p-values are shown; full statistical analyses available in Tables 8a-c. Three biological replicates. Statistical significance was determined at *p* < 0.05. Graphs display mean ± s.e.m.

IL-2 levels in untransduced and Don-CAR-expressing DO11.10 cells remained below the detection threshold (orange line; Fig. 6b), showing that Aβp3-42 treatments did not engage the activation of the cell clones and were therefore not analyzed. The statistical analysis for E2814-CAR revealed statistically significant differences (*F*(2,6) = 16.5955, *p* = 0.0036, *η²* = 0.8469; Fig 6c; Table 8a). However, multiple comparisons showed there was only a significant reduction in IL-2 levels from 1 to 20 μM (*p* = 0.0031), indicating a lack of response of the CAR to Aβp3-42. In contrast, Rem-CAR exhibited statistically significant effects (*F*(2,6) = 49.3965, *p* = 0.0002, *η²* = 0.9427), driven by a marked increase in IL-2 production above VEH at both of the tested Aβp3-42 concentrations (Fig. 6d; Table 8b). Finally, IL-2 levels in Lec-CAR-expressing cells did not show significant changes across Aβp3-42 treatments (*F*(2,6) = 0.2018, *p* = 0.8226, *η²* = 0.0630; Fig. 6e; Table 8c), indicating a lack of response to Aβp3-42 exposure.

## Discussion

### Advantages of using DO11.10 cells for CAR studies in mouse models of Alzheimer’s Disease

This study evaluated CAR function in mouse DO11.10 cells which is, to the best of our knowledge, a novel strategy to assess mouse CARs. For cancer research, CARs typically utilize human sequences as these are primarily tested in human cell-based interventions targeting human xenografts in immunodeficient mouse models^38^. However, most mouse models of AD are immunocompetent and will consequently reject human cell interventions, necessitating the use of syngeneic mouse immune cell interventions. Testing CARs in primary mouse T cells, however, requires the euthanasia of donor animals, and these murine cells are notoriously difficult to transduce and culture.

DO11.10 cells provide a practical alternative for *in vitro* CAR testing prior to implementation of interventions with primary mouse cells. These DO11.10 cells express a monoclonal TCR specific for OVA^22^, which prevents unintended TCR activation by tau and Aβ aggregates while enabling direct evaluation of CAR activation via conserved CD3ζ signaling. Besides their functional advantages, DO11.10 cells are robust, easy to culture, and align with ethical research practices by eliminating the need for animal use. These characteristics make them highly reproducible and suitable for bridging *in vitro* CAR testing with *in vivo* applications in existing mouse models of AD.

### Assessment of CAR targeting in an AD-like cytotoxic environment

In our experiments, CARs exhibited basal tonic ligand-independent activation, a well-documented feature of CAR constructs^28^. However, we detected additional factors influencing CAR activation. Most CAR applications target membrane-bound ligands on cell surfaces, sparing the CAR-expressing cells from exposure to complex toxic environments that may significantly alter their activation dynamics. In contrast, CARs designed to target tau and Aβ protein aggregates in AD introduce an additional layer of complexity due to the cytotoxic nature of these aggregates, which, as demonstrated in our study, influences the activation dynamics of DO11.10 cells. In tau PFF treatments, the primary effect was a nonspecific suppression of tonic CAR activation. As a result, CAR specificity for tau PFFs became evident only when the activation was strong enough to overcome this suppressive effect. Aβ_1-42_ treatments introduced greater complexity, characterized by both contraction and expansion of cell populations as well as nonspecific activation, likely related to widespread cytotoxicity. Thus, exposure to Aβ_1-42_ could both increase and decrease activation metrics nonspecifically, creating a more intricate interplay in the assessment of target-specific CAR activation.

### CAR Response to Tau PFFs, Aβ***_1-42_***, and Aβp3-42

In this framework, the tau-targeting E2814-CAR showed robust specific response to high tau PFF concentrations across all metrics. In contrast, neither the Aβ-targeting Lec-CAR nor untransduced cells were responsive. For Aβ_1-42_ experiments, Aβ-targeting Adu-CAR and Lec-CAR demonstrated activation. However, Lec-CAR activation was within the range of E2814-CAR response by some metrics, questioning its practical use as an intervention. In contrast, Adu-CAR response was qualitatively superior to all other CAR-expressing clones across all metrics. Our finding that Adu-CAR can detect Aβ_1-42_ aligns with the functionality of previously reported Aducanumab-based CAR-Ms^8^ and of the Adu scFv previously used by us in synNotch receptors^17^. Further, this replication of a functional Adu-CAR confirms the validity of our DO11.10 testing platform, demonstrating consistency in reported functionality observed in other cell types. For Aβp3-42 experiments, only the Aβp3-42-targeting Rem-CAR significantly increased IL-2 levels, highlighting its specificity, while untransduced cells, the Aβp3-42-targeting Don-CAR, the Aβ-targeting Lec-CAR, and the tau PFFs-targeting E2814-CAR failed to respond.

#### Limitations and Future Directions

This study provides significant insights into CAR constructs targeting tau, Aβ, and Aβp3-42 but acknowledges several limitations. Expanding the tau PFF concentration range may better define E2814-CAR activation, although high concentrations (500 nM) already surpassed detection limits of the IL-2 ELISA kit (data not shown). Additional Aβ preparations enriched in protofibrils should be used to fully characterize Lec-CAR selectivity. Future studies should explore alternative CAR architectures^11^ for Lec-CAR and Don-CAR to improve activation responses or discard them as nonfunctional CARs. Additionally, CD69 expression analysis for Rem-CAR would validate activation mechanisms beyond IL-2 secretion.

## Conclusions

Our findings demonstrate that DO11.10 cells provide a reliable syngeneic platform for testing various scFv-based antigen-binding domains in mouse CARs. This approach demonstrates that a large variety of available antibodies with well-characterized targets^1,19^ can be effectively leveraged to design scFvs that selectively target distinct pathological features of AD. Since the effector function of scFvs depends on the type of receptor in which they are incorporated and the type of cell in which they are expressed^8,10–12,17^. Consequently, this model offers an effective platform for *in vitro* evaluation of scFvs in the context of various cell-based interventions.

## Acknowledgments

Funding was provided by NIH RF1 AG068296 & R01 AG081989 (J.K.A., C.C.W., S.L., N.J.B.), and the Diana Jacobs Kalman/AFAR Scholarships for Research in the Biology of Aging (C.J.S.). The DO11.10 cell line was a kind gift from Dr. Herb Kasler. The figures were prepared using BioRender.

## Author Contributions

C.J.S. and C.C.W. conceptualized the study and designed the constructs. N.J.B. performed the in-house subcloning. N.J.B. and S.L. produced the gamma-retroviruses. C.J.S. generated the cell clones and designed the experiments. C.J.S., N.J.B., and S.L. conducted the experiments. C.J.S. processed the data for statistical analysis. C.J.S. and C.C.W. performed the statistical analyses. J.K.A. and C.C.W. supervised and guided the project. C.J.S. and C.C.W. wrote the manuscript, with input from all other authors.

## Declaration of Interests

J.K.A., C.C.W., and S.L. are inventors in Patent No. WO2024076500A3.

## Methods

### Constructs

E2814-CAR (Table 1b), Lec-CAR (Table 1c), Adu-CAR (Table 1d), Don-CAR (Table 1e), and Rem-CAR (Table 1f) are based on a second-generation CAR architecture featuring a mouse CD28 costimulatory domain (Fig. 1a; Table 1a), based on the 1D3-28Z sequence (Kochenderfer et al., 2010). All CAR constructs were directed to the cell membrane using a human CD8α signal sequence, with their scFv sequences comprising the VLC positioned at the 5’ end of the VHC. Each CAR construct was expressed upstream of a mFoxp3 gene, separated by a T2A sequence (Fig. 1b). Downstream of the mFoxp3 gene, an EGFP reporter gene was separated using a P2A sequence. The inclusion of mFoxp3 in the expression cassette aimed to induce a Treg phenotype in mouse CD4+ T cells^39,40^ as part of a CAR-Treg intervention strategy under development in our laboratory that was not assessed by this study. Both 2A sequences included a Furin cleavage site and a GSG linker to minimize C-terminal amino acid additions caused by 2A ribosomal skipping^41^. The mouse codon-optimized Adu-CAR sequences were synthesized by Genescript and subcloned into the MIGR1 transfer plasmid by PCR. MIGR1 was a kind gift from Dr. Warren Pear (Addgene plasmid # 27490; http://n2t.net/addgene:27490; RRID:Addgene_27490). The MIGR1 IRES was replaced with T2A, mFoxp3, and P2A sequences to enable strong multicistronic expression of the CAR, mFoxp3, and EGFP (Fig. 1b). Mouse codon-optimized T2A, P2A, and mFoxp3, were synthesized by Vector Builder. Lec-CAR was built by using the scFv from the mouse codon-optimized chimeric human-mouse Lecanemab scFv-Fc (chLecanemab) antibody previously described by us^17^. Briefly, the scFv from chLecanemab was substituted by PCR for the scFv of Adu-CAR in the multicistronic MIGR1 expression cassette to create Lec-CAR MIGR1 (Fig. 1b). The sequences for E2814-CAR, Don-CAR, and Rem-CAR in the same multicistronic MIGR1 expression cassette sequence (Fig. 1b) were codon-optimized for mice, synthesized, and subcloned by Vector Builder. All constructs were sequence-verified by Genescript, Vector Builder and or Primordium Labs. In-house subcloning was performed by Gibson Assembly using the NEBuilder® HiFi DNA Assembly Cloning Kit (NEB, CAT# E5520S) and Q5 Hot Start High-Fidelity 2X Master Mix (NEB, CAT# M0494L).

### Gamma retrovirus production

Platinum-E (Plat-E) (Cell Biolabs, CAT# RV-101) cells were used for viral production. Following the manufacturer’s instructions, retroviral transfer plasmids were transfected into Plat-E cells using Lipofectamine™ 3000 Transfection Reagent (Invitrogen, CAT# L3000001). Retroviral supernatants were harvested 2 days post-transfection, centrifuged at 800 x g for 10 minutes, filtered through a 0.45 µm filter (VWR, CAT#76479-020), and subsequently frozen at −80°C for long-term storage.

### Cell culture, transduction, and clone purification

#### Cell culture

The DO11.10 murine hybridoma T-cell line, generously provided by Dr. Herb Kasler, was cultured in suspension using non-treated plates or flasks at 37°C in a humidified atmosphere with 5% CO₂. Cells were cultured in complete culture media comprising Roswell Park Memorial Institute (RPMI) 1640 Medium with GlutaMAX™ Supplement (Gibco, CAT# 61870-036), 10% Fetal Bovine Serum (FBS; Corning, CAT# 35-010-CV), 1 mM sodium pyruvate (Gibco, CAT# 11360-070), 100 U/mL penicillin and 100 µg/mL streptomycin (1X Pen/Strep; Gibco, CAT# 15140122), and 1X 2-mercaptoethanol (Gibco, CAT# 21985-023). The medium was vacuum filtered using a GenClone Vacuum Filter System with a PES Membrane (0.22 µm; Genesee Scientific, CAT# 25-227). **Transduction.** Cells in transduction media (complete culture media lacking Pen/Strep) were seeded at a density of 1 × 10⁵ cells per well in untreated 12-well plates. Each well received polybrene transduction enhancer (**VectorBuilder, CAT# PL0001**) diluted in transduction media and in-house-produced viral particle preparations, resulting in each well containing 2 mL of 1 × 10⁵ cells with 5 µg/mL polybrene and 1 mL of viral particle preparations. The plates were incubated at 37°C with 5% CO₂ for 48 hours. After this incubation period, the viral particle-containing media was replaced with fresh complete culture media. **Clone Purification**. Twenty-four hours after replacing the viral media, cells were washed with Dulbecco’s Phosphate Buffered Saline 1X (DPBS; VWR, CAT# 02-01119-0500) and stained with the LIVE/DEAD™ Fixable Violet Dead Cell Stain Kit (ThermoFisher, CAT# L34964) at room temperature for 20 minutes. Cells were then resuspended in FACS buffer (DPBS with 1% FBS and 10 mM HEPES; Gibco, CAT# 15630080). Transduced cells were sorted based on EGFP expression using a BD FACSAria II cell sorter (BD Biosciences). Transduced clones were expanded and subjected to a second round of FACS. In this step, cells were sorted based on a range of EGFP expression to ensure comparable copy-number integration of the expression cassette across CAR clones (Fig. 1d). Cells were cryopreserved in 90% FBS and 10% DMSO.

### Peptide preparation and treatments

#### Preparation

Tau-441 preformed fibrils (Tau PFFs; rPeptide, CAT# TF-1001-2) were purchased as a 1 mg/mL stock solution. For Tau PFF experiments, DPBS was used as the vehicle control. Beta-amyloid (1-42) (Aβ_1-42_; rPeptide, CAT# A-1167-2) and [Pyr3]-beta-Amyloid (3-42) (Aβp3-42; Anaspec, CAT# AS-29907) were aggregated using the Beta Amyloid (1-42) Aggregation Kit (rPeptide, CAT# A-1170) following the manufacturer’s protocol. Briefly, lyophilized Aβ_1-42_ and Aβp3-42 peptides were dissolved in 5 mM Tris or 10 mM NaOH at a concentration of 1 mg/ml. These solutions were then diluted with HPLC water and Tris-buffered saline (TBS) to create 100 µM solutions. These solutions were incubated for 3 hours at 37°C to produce 100 µM stock solutions enriched in Aβ_1-42_ oligomers or Aβp3-42 aggregates. The vehicle stock solution used for Aβ_1-42_ and Aβp3-42 experiments consisted of 5 mM Tris or 10 mM NaOH, diluted in HPLC water and TBS in the same ratio as the amyloid stock solutions, and was similarly incubated for 3 hours at 37°C. All peptide stock solutions were serially diluted with complete culture media to obtain the desired treatment concentrations. **Treatments.** Cells were seeded in suspension in non-treated well plates at a final density of 1 × 10⁵ cells/mL. For treatments with Tau PFFs, cells were incubated with Tau PFFs or the vehicle for 45 hours at 37°C in a humidified atmosphere with 5% CO₂ prior to harvest. For treatments with Aβ_1-42_ or Aβp3-42 aggregates, cells were exposed to these peptides or the respective vehicles for 48–49 hours under the same conditions. All treatment experiments were performed in triplicate.

### IL-2 ELISA

The OptEIA™ Mouse IL-2 ELISA Set (BD, CAT# 555148) was used to assess IL-2 production by DO11.10 cells according to manufacturer’s instructions, as a measure of T-cell activation (Roy 2012). Briefly, microwells of the assay plate were coated with 100 µL per well of anti-mouse IL-2 capture antibody (BD, CAT# 51-26141E) diluted 1:250 in coating buffer (0.1 M Sodium Carbonate, pH 9.5), followed by overnight incubation at 4°C. Wells were washed three times with 300 µL/well wash buffer (0.05% Tween-20 in DPBS). Plates were blocked with 200 µL/well assay diluent (10% FBS in DPBS) and incubated at room temperature (RT) for 1 hour, followed by washing as described. Standards were made with recombinant mouse IL-2 (BD, CAT# 51-26146E) and serially diluted with assay diluent. Then, 100 µL of each standard or undiluted sample supernatant was transferred to its designated well on the assay plate. The plate was sealed and incubated for 2 hours at RT. Wells were washed five times. Next, 100 µL of working detector solution, containing 1:250 Biotinylated anti-mouse IL-2 detection antibody (BD, CAT# 51-26142E) and 1:250 Streptavidin-horseradish peroxidase enzyme reagent (BD, CAT#51-9002813) in assay diluent, was added to each well. The assay plate was sealed and incubated for 1 hour at RT. Wells were washed seven times. Subsequently, 100 µL of tetramethylbenzidine substrate solution (BD, CAT# 555214) was added to each well, and the plate was incubated for 30 minutes at RT in the dark. Finally, 50 µL of Stop Solution (1 M H_3_PO_4_) was added to each well, and absorbance was read immediately at 450 nm, with wavelength correction at 570 nm, using the BioTek Cytation 3 (Agilent). The supernatant of each sample was tested for IL-2 in technical triplicates. ELISA data was analyzed using GainData (Arigo Biolaboratories, https://www.arigobio.com/elisa-analysis). Standards were plotted using Four parameter logistic (4PL) modeling to perform IL-2 quantification of each sample in pg/mL.

### Flow cytometry

After treatment, cells were pelleted and washed with DPBS for flow cytometry analysis. The cells were stained with a fixable viability dye and antibodies, diluted in DPBS, for 45 min at 4°C. The following viability dye and antibodies were used: LIVE/DEAD™ Fixable Violet Dead Cell Stain Kit (1:1000, Invitrogen, CAT# L34964), Anti-mouse CD69 APC-eFluor™ 780 (1:80, H1.2F3, eBioscience, CAT# 47-0691-82), and Anti-mouse CD69-PeCy7 (1:200, H1.2F3, BioLegend, CAT# 104512). After staining, cells were washed in DPBS and resuspended in FACS buffer containing 1% FBS and 10mM HEPES in DPBS. The flow cytometer Cytek Aurora (Cytek) was used to measure viability, the transduction reporter EGFP, and CD69. Gating was based on fluorescence-minus-one (FMO) controls included for each marker.

### Statistical analysis

Statistical analyses were conducted using IBM SPSS Statistics (RRID: SCR_002865) and the built-in statistical analysis tools in Biorender (Biorender.com) powered by R (RRID: SCR_001905). Only statistically significant results or notable non-significant trends are reported in the text and reflected in the figure graphs. A complete set of analyses and results is provided in Tables 2–8. Assumption checks were conducted before performing parametric tests. Outliers were identified using box plots, normality was assessed using the Shapiro-Wilk test (p > 0.05) and Q-Q plots, and homogeneity of variances was evaluated using Levene’s test based on means (p > 0.05) and scatter plots of predicted versus residual values. When possible, true value outliers were substituted by the closest rank value plus or minus 1% of the range of the datapoints and the assumptions tests re-run. Groups of true value outliers reflecting a strong response to an intervention were not modified. All outliers and substitutions are noted in the supplementary tables. If normality and homogeneity of variances assumptions were not met, the data was transformed appropriately to satisfy the requirements and tests re-run, or alternative Welch’s F test with Games-Howell multiple comparisons, or independent t-test for lack of homogeneity of variances, or non-parametric Kruskal-Wallis test with Dunn’s multiple comparisons were employed as required. When assumptions were met, independent t-tests, one-way or two-way ANOVAs were performed as needed, the latter followed by Dunnett or Tukey’s tests for comparisons to the VEH or all pairwise comparisons, respectively. **Viability.** The total number of live untransduced and CAR-expressing clones (EGFP- and EGFP+) were analyzed to assess viability. Multiple comparisons correcting the familywise error were made between each treatment condition and the vehicle control. **CD69**. The percentage of CD69-positive cells was used to assess activation. All pairwise comparisons among all treatment conditions were performed to interpret significant omnibus test results. **IL-2.** For ELISA assays, the lowest measurable IL-2 concentration reported for the OptEIA™ Mouse IL-2 ELISA kit was 3.1 pg/mL. To mitigate the impact of missing data on statistical analyses, a constant value of 3 was added to all data points, and the resulting values were log10-transformed. To avoid excessive familywise error correction, only meaningful treatment conditions with IL-2 values exceeding those of vehicle controls were included in the comparisons. CAR constructs that showed no IL-2 levels above vehicle control in all treatment conditions were excluded from the analysis. **IL-2 signal-to-noise ratio.** For Aβ_1-42_ experiments, the signal-to-noise ratio was calculated to compare CAR constructs without the confounding effects of tonic ligand-independent activation. The signal-to-noise ratio was determined by vehicle normalization, wherein the fold increase in IL-2 for each treatment condition was calculated relative to the vehicle control for each CAR construct. To meet parametric test assumptions, vehicle-normalized vehicle control conditions with zero variance were excluded from the analysis. To meet the assumption of homogeneity of variances of the two-way ANOVA, the values were square root-transformed to address violations caused by residuals increasing with predicted values. All pairwise comparisons were two-tailed. Exact p-values are reported for significant and non-significant results.

## References

1. Cummings, J., Osse, A. M. L., Cammann, D., Powell, J. & Chen, J. Anti-Amyloid Monoclonal Antibodies for the Treatment of Alzheimer’s Disease. BioDrugs 38, 5–22 (2024).

2. Hyman, B. T. et al. National Institute on Aging–Alzheimer’s Association guidelines for the neuropathologic assessment of Alzheimer’s disease. Alzheimers Dement. 8, 1–13 (2012).

3. DeTure, M. A. & Dickson, D. W. The neuropathological diagnosis of Alzheimer’s disease. Mol. Neurodegener. 14, 32 (2019).

4. Jack, C. R. et al. NIA-AA Research Framework: Toward a biological definition of Alzheimer’s disease. ALZHEIMERS Dement. 14, 535–562 (2018).

5. Cummings, J. et al. Alzheimer’s disease drug development pipeline: 2024. Alzheimers Dement. Transl. Res. Clin. Interv. 10, e12465 (2024).

6. Saetzler, V. et al. Development of beta-amyloid-specific CAR-Tregs for the treatment of Alzheimer’s disease. Cells 12, 2115 (2023).

7. Yeapuri, P. et al. Amyloid-β specific regulatory T cells attenuate Alzheimer’s disease pathobiology in APP/PS1 mice. Mol Neurodegener 18, 97 (2023).

8. Kim, A. B., et al. Chimeric antigen receptor macrophages target and resorb amyloid plaques. JCI Insight 9, (2024).

9. June, C. H. & Sadelain, M. Chimeric antigen receptor therapy. N Engl J Med 379, 64–73 (2018).

10. Manhas, J., Edelstein, H. I., Leonard, J. N. & Morsut, L. The evolution of synthetic receptor systems. Nat. Chem. Biol. 18, 244–255 (2022).

11. Rafiq, S., Hackett, C. S. & Brentjens, R. J. Engineering strategies to overcome the current roadblocks in CAR T cell therapy. Nat Rev Clin Oncol 17, 147–167 (2020).

12. Ferreira, L. M. R., Muller, Y. D., Bluestone, J. A. & Tang, Q. Next-generation regulatory T cell therapy. Nat. Rev. Drug Discov. 18, 749–769 (2019).

13. Maude, S. L. et al. Tisagenlecleucel in children and young adults with B-cell lymphoblastic leukemia. N Engl J Med 378, 439–448 (2018).

14. Neelapu, S. S. et al. Axicabtagene ciloleucel CAR T-cell therapy in refractory large B-cell lymphoma. N Engl J Med 377, 2531–2544 (2017).

15. Morsut, L. et al. Engineering Customized Cell Sensing and Response Behaviors Using Synthetic Notch Receptors. Cell 164, 780–791 (2016).

16. Roybal, K. T. et al. Engineering T Cells with Customized Therapeutic Response Programs Using Synthetic Notch Receptors. Cell 167, 419–432.e16 (2016).

17. Bergo, N. J., Lee, S., Siebrand, C. J., Andersen, J. K. & Walton, C. C. Aβ-targeting synNotch Receptor for Alzheimer’s Disease: Expanding Applications to Extracellular Protein Aggregates. 2024.10.15.618096 Preprint at 10.1101/2024.10.15.618096 (2024).

18. Simic, M. S. et al. Programming tissue-sensing T cells that deliver therapies to the brain. Science 386, eadl4237 (2024).

19. Song, C. et al. Immunotherapy for Alzheimer’s disease: targeting β-amyloid and beyond. Transl. Neurodegener. 11, 18 (2022).

20. Braak, H., Alafuzoff, I., Arzberger, T., Kretzschmar, H. & Del Tredici, K. Staging of Alzheimer disease-associated neurofibrillary pathology using paraffin sections and immunocytochemistry. Acta Neuropathol 112, 389–404 (2006).

21. Braak, H. & Braak, E. Neuropathological stageing of Alzheimer-related changes. Acta Neuropathol 82, 239–259 (1991).

22. Robertson, J. M., Jensen, P. E. & Evavold, B. D. DO11.10 and OT-II T Cells Recognize a C-Terminal Ovalbumin 323–339 Epitope1. J. Immunol. 164, 4706–4712 (2000).

23. Kochenderfer, J. N., Yu, Z., Frasheri, D., Restifo, N. P. & Rosenberg, S. A. Adoptive transfer of syngeneic T cells transduced with a chimeric antigen receptor that recognizes murine CD19 can eradicate lymphoma and normal B cells. Blood 116, 3875–3886 (2010).

24. Nguyen, P., Moisini, I. & Geiger, T. L. Identification of a murine CD28 dileucine motif that suppresses single-chain chimeric T-cell receptor expression and function. Blood 102, 4320–4325 (2003).

25. Roberts, M. et al. Pre-clinical characterisation of E2814, a high-affinity antibody targeting the microtubule-binding repeat domain of tau for passive immunotherapy in Alzheimer’s disease. Acta Neuropathol. Commun. 8, 13 (2020).

26. Cibrián, D. & Sánchez-Madrid, F. CD69: from activation marker to metabolic gatekeeper. Eur J Immunol 47, 946–953 (2017).

27. Boyman, O. & Sprent, J. The role of interleukin-2 during homeostasis and activation of the immune system. Nat Rev Immunol 12, 180–190 (2012).

28. Ajina, A. & Maher, J. Strategies to Address Chimeric Antigen Receptor Tonic Signaling. Mol. Cancer Ther. 17, 1795–1815 (2018).

29. Englund, H. et al. Sensitive ELISA detection of amyloid-beta protofibrils in biological samples. J Neurochem 103, 334–345 (2007).

30. Lord, A. et al. An amyloid-β protofibril-selective antibody prevents amyloid formation in a mouse model of Alzheimer’s disease. Neurobiol. Dis. 36, 425–434 (2009).

31. Sevigny, J. et al. The antibody aducanumab reduces Aβ plaques in Alzheimer’s disease. Nature 537, 50– 56 (2016).

32. Söderberg, L. et al. Lecanemab, Aducanumab, and Gantenerumab — Binding Profiles to Different Forms of Amyloid-Beta Might Explain Efficacy and Side Effects in Clinical Trials for Alzheimer’s Disease. Neurotherapeutics 20, 195–206 (2023).

33. Tucker, S. et al. The Murine Version of BAN2401 (mAb158) Selectively Reduces Amyloid-β Protofibrils in Brain and Cerebrospinal Fluid of tg-ArcSwe Mice. J. Alzheimers Dis. 43, 575–588 (2015).

34. Cline, E. N., Bicca, M. A., Viola, K. L. & Klein, W. L. The amyloid-β oligomer hypothesis: beginning of the third decade. J. Alzheimers Dis. 64, 567–610 (2018).

35. Jarosz-Griffiths, H. H., Noble, E., Rushworth, J. V. & Hooper, N. M. Amyloid-β receptors: The Good, the bad, and the prion protein. J Biol Chem 291, 3174–3183 (2016).

36. Demattos, R. B. et al. A plaque-specific antibody clears existing β-amyloid plaques in Alzheimer’s disease mice. Neuron 76, 908–920 (2012).

37. Mintun, M. A. et al. Donanemab in Early Alzheimer’s Disease. N. Engl. J. Med. 384, 1691–1704 (2021).

38. Siegler, E. L. & Wang, P. Preclinical Models in Chimeric Antigen Receptor–Engineered T-Cell Therapy. Hum. Gene Ther. 29, 534–546 (2018).

39. Fontenot, J. D., Gavin, M. A. & Rudensky, A. Y. Foxp3 programs the development and function of CD4+CD25+ regulatory T cells. Nat Immunol 4, 330–336 (2003).

40. Hori, S., Nomura, T. & Sakaguchi, S. Control of Regulatory T Cell Development by the Transcription Factor Foxp3. Science 299, 1057–1061 (2003).

41. Yang, S. et al. Development of optimal bicistronic lentiviral vectors facilitates high-level TCR gene expression and robust tumor cell recognition. Gene Ther. 15, 1411–1423 (2008).

