## Supplementary Materials for "Chimeric Antigen Receptors Discriminate Between Tau and Distinct Amyloid-Beta Species"

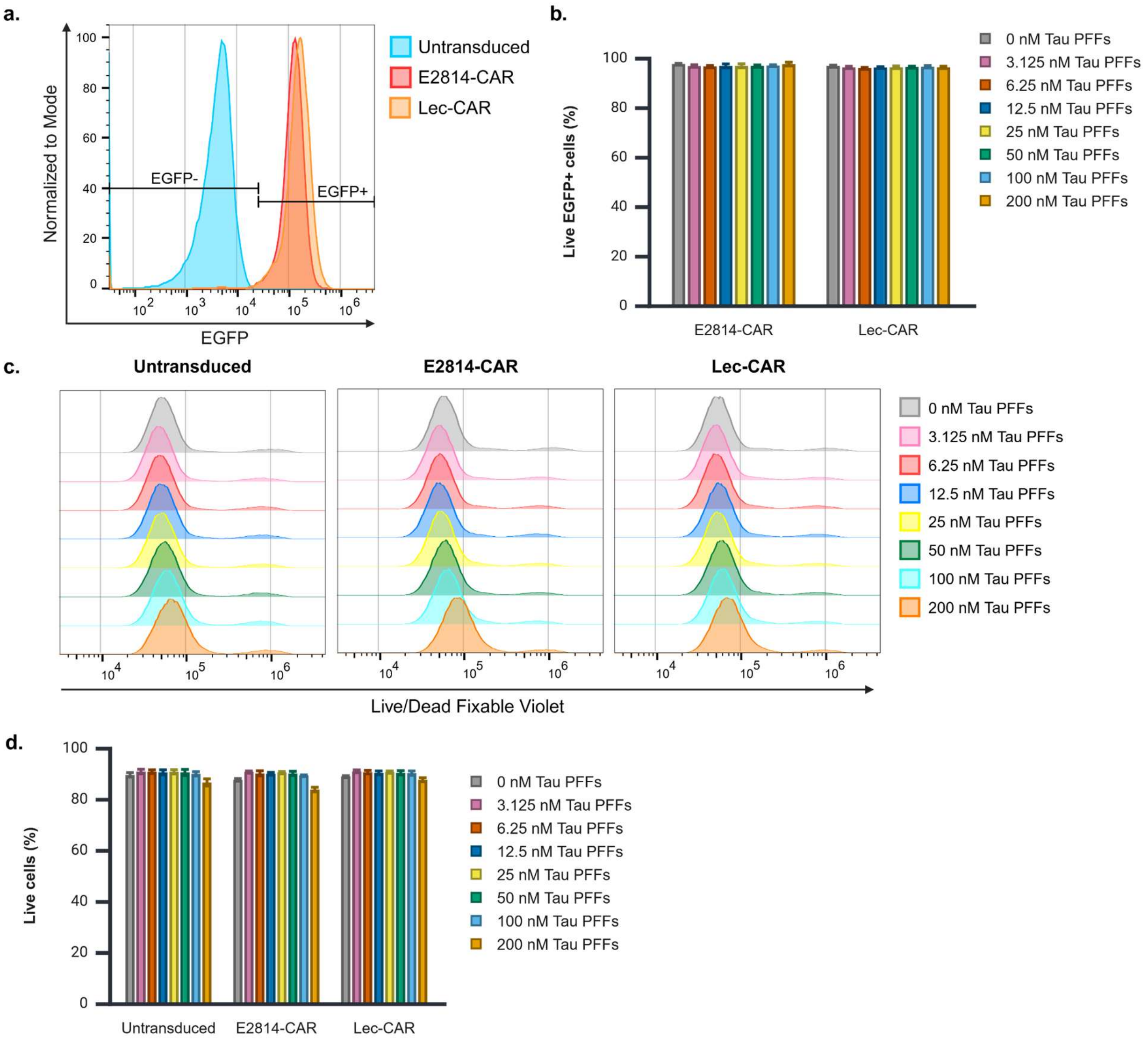


**Supplementary Figure 1. EGFP-expression and viability in tau PFFs experiments.** (**a**) Representative flow cytometry histogram showing negative EGFP signal in untransduced cells and positive EGFP signal in E2814-CAR and Lec-CAR cell clones under VEH treatment conditions. (**b**) Mean percentage of live EGFP+ E2814-CAR and Lec-CAR cell clones across all experiments. (**c**) Representative flow cytometry histograms of live/dead fixable violet staining for untransduced, E2814-CAR, and Lec-CAR clones under tau PFF treatment conditions, demonstrating comparable viability. (**d**) Mean percentage of live cells, including untransduced (EGFP-) and CAR-expressing clones (EGFP+ and EGFP-), across all experiments. 3 biological replicates. Bar graphs display mean ± S.D.

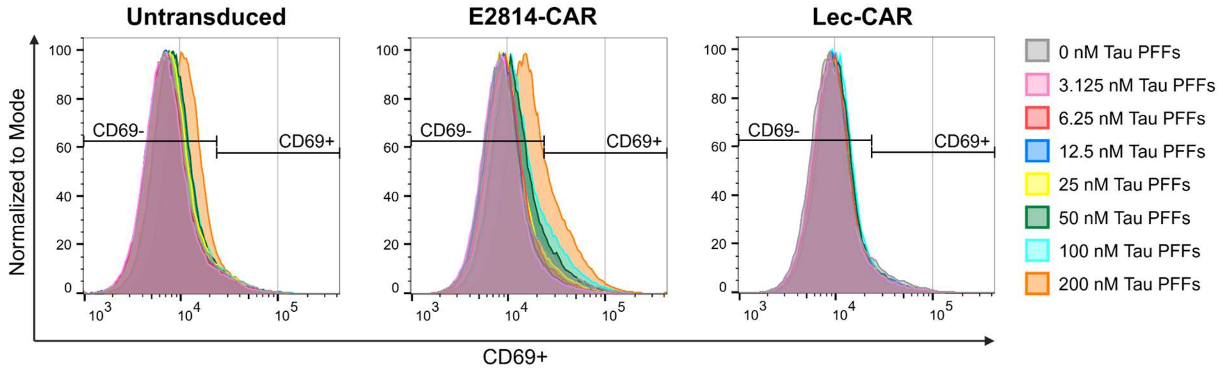


**Supplementary Figure 2. CD69-expression for tau PFF experiments.** Representative flow cytometry histograms of CD69 signal for untransduced cells, E2814-CAR clones, and Lec-CAR clones under tau PFF treatment conditions.

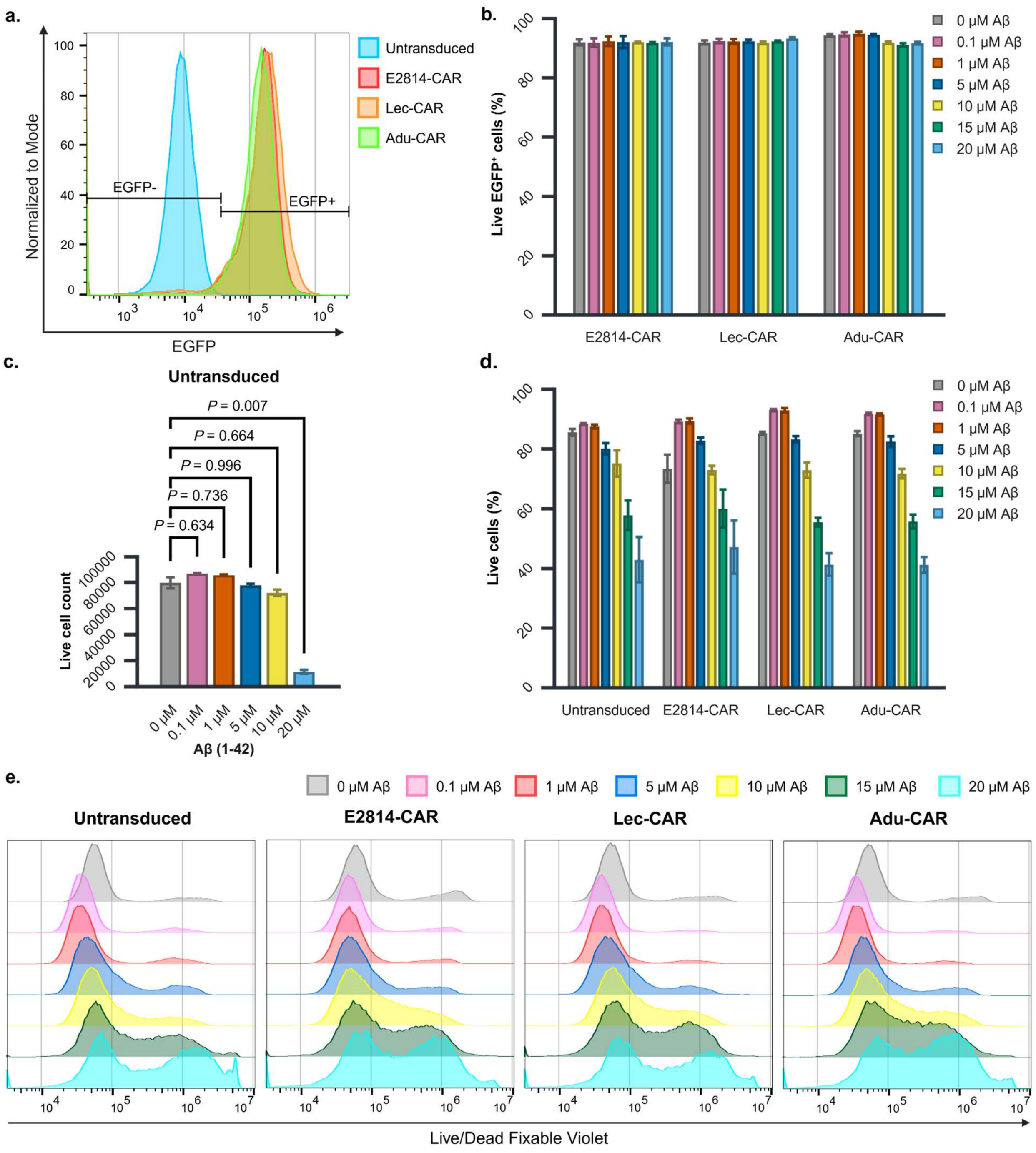


**Supplementary Figure 3. EGFP-expression and viability in Aβ$_{1-42}$ experiments.** (**a**) Representative flow cytometry histogram showing negative EGFP signal in untransduced cells and positive EGFP signal in E2814-CAR, Lec-CAR cells, and Adu-CAR cell clones under VEH treatment conditions. (**b**) Mean percentage of live EGFP+ E2814-CAR, Lec-CAR, and Adu-CAR cell clones across all experiments. 3 biological replicates. Bar graphs display mean ± S.D. (**c**) Welch's one-way ANOVA with Games-Howell multiple comparison of live cell count of untransduced cells excluding the non-normally distributed 15 μM Aβ$_{1-42}$ treatment group. 3 biological replicates. Statistical significance was determined at $p < 0.05$. Graphs display mean ± S.E.M. (**d**) Mean percentage of live cells, including untransduced (EGFP-) and CAR-expressing clones (EGFP+ and EGFP-), across all experiments. 3 biological replicates. Bar graphs display mean ± S.D. (**e**) Representative flow cytometry histograms of live/dead fixable violet staining for untransduced, E2814-CAR, Lec-CAR, and Adu-CAR clones at indicated Aβ$_{1-42}$ concentrations, demonstrating increased cell death in higher Aβ$_{1-42}$ concentrations.

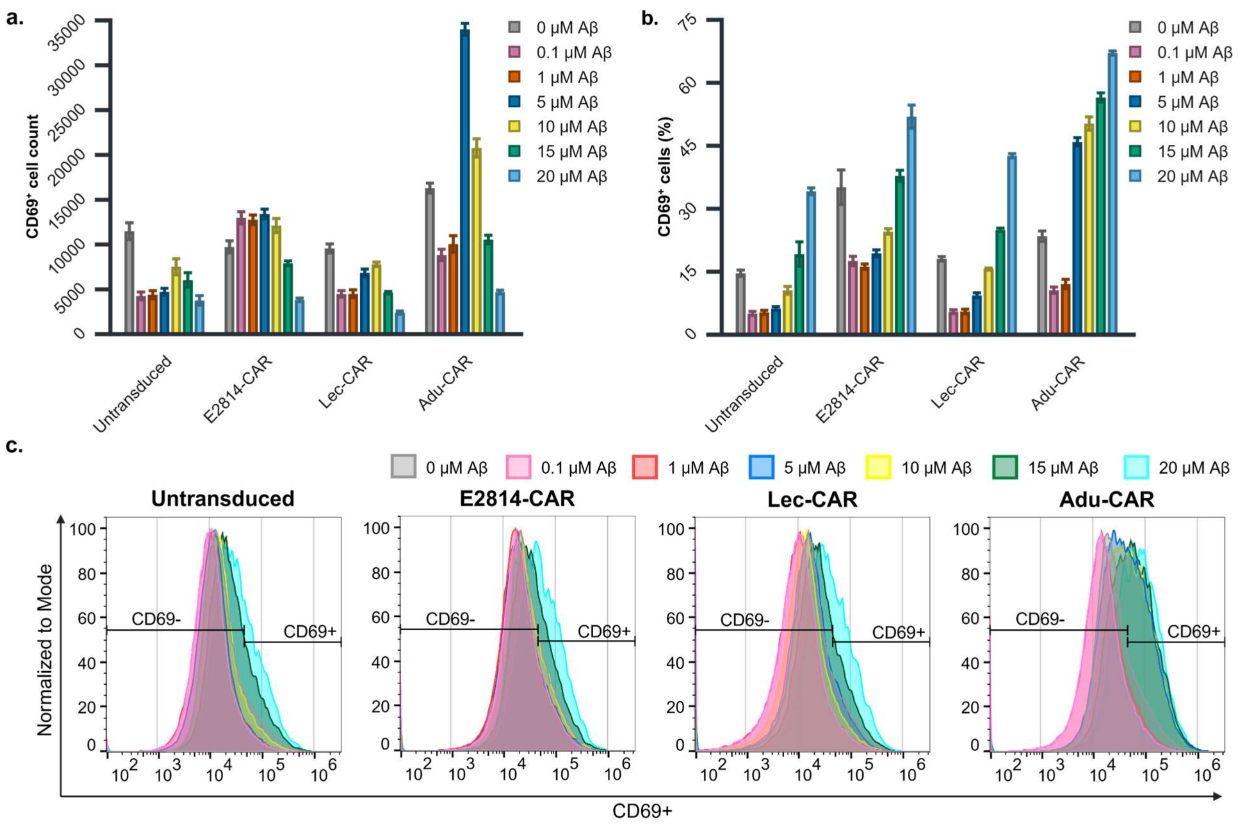


**Supplementary Figure 4. CD69-expression in A$\beta_{1\text{-}42}$ experiments.** (**a**) Mean total count of CD69+ cells and (**b**) percentage of CD69+ cells for untransduced, E2814-CAR, Lec-CAR, and Adu-CAR groups across indicated A$\beta_{1\text{-}42}$ concentrations for all experiments. The reduction in total CD69+ cell count in (**a**) corresponds with decreased overall viability observed in Supplementary Figure 3d, while the percentage of activated CD69+ cells in (**b**) increases with reduced viability. 3 biological replicates. Bar graphs display mean ± SD. (**c**) Representative flow cytometry histograms of CD69 signal for untransduced cells, E2814-CAR clones, Lec-CAR clones, and Adu-CAR clones at indicated A$\beta_{1\text{-}42}$ concentrations.

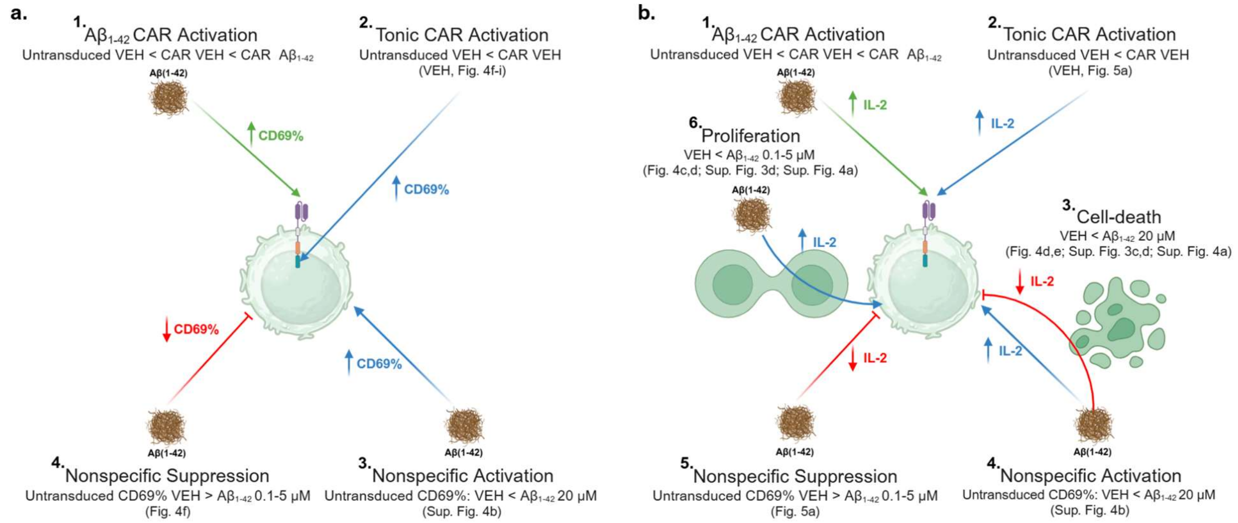


**Supplementary Figure 5. CAR-specific and nonspecific effects of $A\beta_{1-42}$ treatments.** Net activation as assessed by CD69 (**a**) and IL-2 (**b**) reflects the combined effects of $A\beta_{1-42}$-specific CAR activation (green), nonspecific activation and or proliferation (blue), and nonspecific suppression and or cell loss (red). Due to the inherent methodological differences, each metric is affected differently. (**a**) Variables contributing to net activation as measured by the percentage of CD69-positive cells. (**1**) CAR-specific ligand-dependent activation: $A\beta_{1-42}$ treatment leading to an increased percentage of CD69-positive cells relative to CAR VEH-treated cells and untransduced controls. (**2**) Tonic ligand-independent activation: Constitutive CAR activation evidenced by a higher percentage of CD69-positive cells in VEH-treated CAR cells (Fig. 4g-i) compared to VEH-treated untransduced cells (Fig. 4f). Different constructs display varying levels of tonic activation, which should proportionally increase the response to $A\beta_{1-42}$. Consequently, the direct comparison between constructs is unreliable without prior normalization to VEH (signal-to-noise ratio). (**3**) Nonspecific activation: At 20 μM $A\beta_{1-42}$ untransduced cells exhibited nonspecific activation (Sup. Fig. 4b). This indicates nonspecific activation is contributing to the response of all CAR clones, likely to affect lower concentrations as well. A potential explanation is a nonspecific response to extensive cell death seen at this concentration (Fig. 4d, e; Sup. Fig. 3c, d; Sup. Fig. 4a). (**4**) Nonspecific suppression: Suppressive effects are observed in both CAR-expressing and untransduced cells at low $A\beta_{1-42}$ concentrations (Fig. 4f-i). In contrast to tau PFF treatments (Fig. 2e), this suppression was also present in untransduced cells (Fig. 4f) and indicated the involvement of alternative and/or stronger suppressive mechanisms. (**b**) Variables contributing to net activation as measured by IL-2 production. In contrast to the percentage of CD69-positive cells, IL-2 levels are directly influenced by the absolute number of activated cells as well as the magnitude of their activation. (**1**) CAR-specific activation is as explained in Sup. Fig. 5a1. (**2**) Tonic CAR activation is as explained in Sup. Fig. 5a2, evidenced in Fig. 5a. (**3**) Cell death: 20 μM $A\beta_{1-42}$ treatments reduced overall cell viability (Fig. 4d, e; Sup. Fig. 3c, d) and the total number of activated CD69-positive cells (Sup. Fig. 4a), shrinking the population of tonically activated CAR clones as well as $A\beta_{1-42}$-responsive CAR clones. (**4**) Nonspecific activation: Detected by the analysis of CD69 in untransduced cells at 20 μM $A\beta_{1-42}$ (Sup. Fig. 4b) can be expected to result in increased IL-2 production. Consequently, at higher $A\beta_{1-42}$ concentrations, there are likely CAR-independent forces driving increases and decreases in IL-2 linked to nonspecific activation (Sup. Fig. 5b.4) and cell loss (Sup. Fig. 5b.3), respectively. (**5**) Nonspecific suppression: Suppression of IL-2 production is seen for all CAR clones in Fig. 5a. (**6**) Proliferation: Expansion of IL-2-producing cells seen at lower $A\beta_{1-42}$ concentrations (Fig. 4c, d; Sup. Fig. 3d; Sup. Fig. 4a) can contribute to increased IL-2 levels, albeit this should be considered in the context of a reduction in the percentage of CD69-positive cells at overlapping concentrations (Sup. 5a.4).
